# Preferential catabolism of L- vs D-serine by *Proteus mirabilis* contributes to pathogenesis and catheter-associated urinary tract infection

**DOI:** 10.1101/2022.06.02.494593

**Authors:** Aimee L. Brauer, Brian S. Learman, Steven M. Taddei, Namrata Deka, Benjamin C. Hunt, Chelsie E. Armbruster

## Abstract

*Proteus mirabilis* is a common cause of urinary tract infection, especially in catheterized individuals. Amino acids are the predominant nutrient for bacteria during growth in urine, and our prior studies identified several amino acid import and catabolism genes as fitness factors for *P. mirabilis* catheter-associated urinary tract infection (CAUTI), particularly D- and L-serine. In this study, we sought to determine the hierarchy of amino acid utilization by *P. mirabilis* and to examine the relative importance of D- vs L-serine catabolism for critical steps in CAUTI development and progression. Herein, we show that *P. mirabilis* preferentially catabolizes L-serine during growth in human urine, followed by D-serine, threonine, tyrosine, glutamine, tryptophan, and phenylalanine. Independently disrupting catabolism of either D- or L-serine has minimal impact on *in vitro* phenotypes while completely disrupting both pathways decreases motility, biofilm formation, and fitness due to perturbation of membrane potential and cell wall biosynthesis. In a mouse model of CAUTI, loss of either serine catabolism system decreased fitness, but disrupting L-serine catabolism caused a greater fitness defect than disrupting D-serine catabolism. We therefore conclude that hierarchical utilization of amino acids may be a critical component of *P. mirabilis* colonization and pathogenesis within the urinary tract.

**Abbreviated Summary:** Amino acids are a predominant nutrient in urine, and their import and catabolism has been hypothesized to contribute to the ability of bacteria to cause urinary tract infection. We demonstrate that a common uropathogen, *Proteus mirabilis,* preferentially catabolizes L-serine followed by D-serine, threonine, tyrosine, and glutamine during growth in human urine. We further demonstrate that L-serine catabolism provides a greater fitness advantage than D-serine catabolism, yet both pathways contribute to pathogenesis in the urinary tract.

**Graphical Abstract:** (Created using BioRender.com)

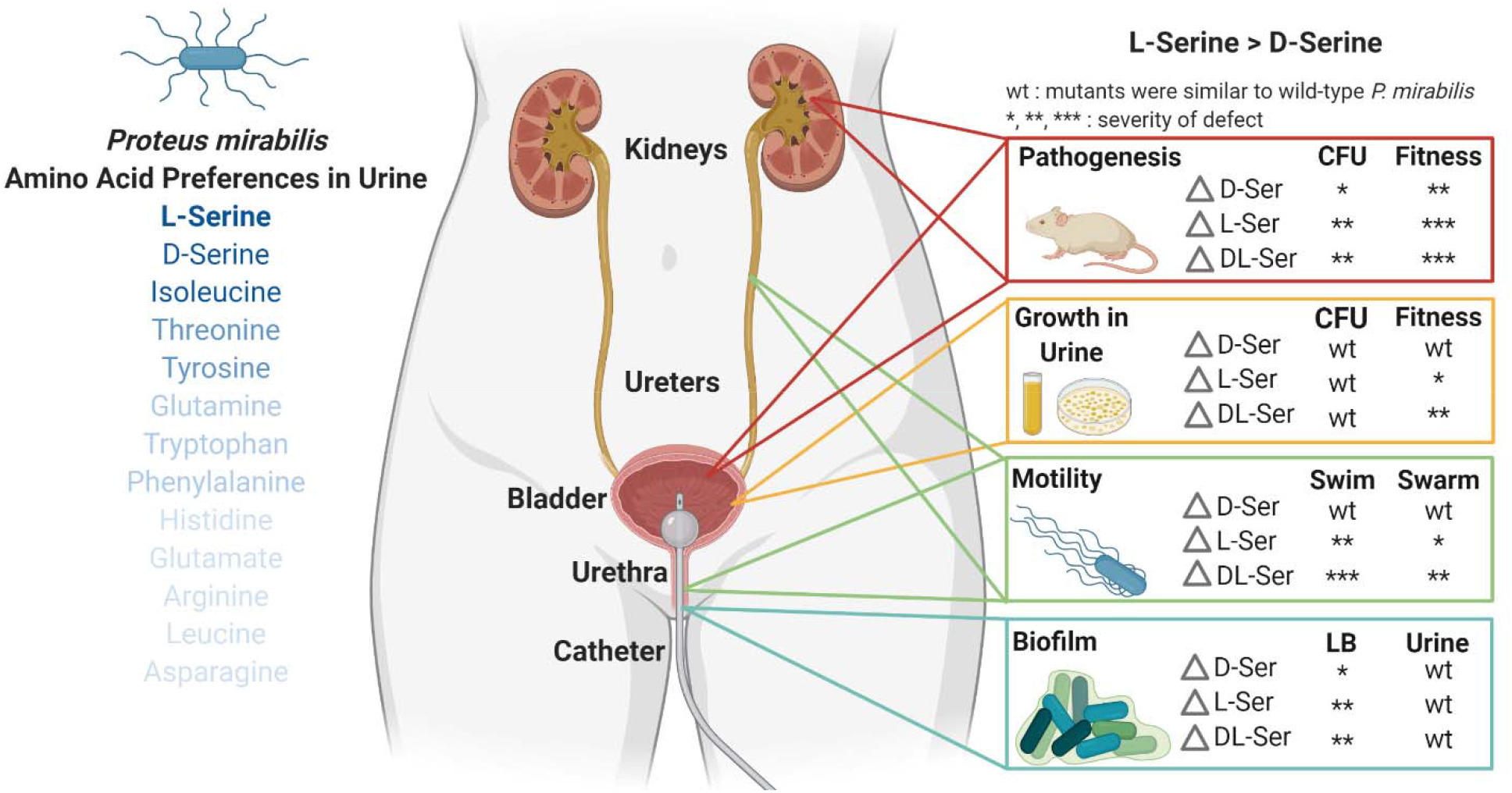

## Introduction

Urinary tract infections (UTIs) are among the most common infections worldwide and a leading cause of morbidity and healthcare expenditures across all ages in the United States. UTIs are broadly classified as being uncomplicated or complicated based on patient risk factors for severe disease and localization of symptoms (Wagenlehner *et al*., 2020, Sabih & Leslie, 2022). Uncomplicated UTI refers to infection in the absence of known risk factors for complications or treatment failure and can be further divided into acute cystitis (lower urinary tract symptoms) or acute pyelonephritis (upper urinary tract symptoms). Complicated UTI refers to infections that carry a high risk of treatment failure, recurrence, or morbidity and mortality, which includes infections that occur in pregnant women, immunocompromised patients, the elderly, and individuals with urinary catheters. The vast majority of uncomplicated UTIs are caused by *Escherichia coli,* while the etiology of complicated infection is more diverse and includes *E. coli, Proteus mirabilis, Klebsiella* spp., *Enterococcus faecalis, Pseudomonas aeruginosa,* and *Staphylococcus* spp. among others (Gaston *et al*., 2021).

In order to establish a UTI, microbes that gain entry to the urinary tract must first adapt to growth in urine. Mounting evidence indicates that a combination of flexible metabolism and the ability to grow rapidly in urine are among the most important indicators of pathogenic potential within the urinary tract (Reitzer & Zimmern, 2019, Forsyth *et al*., 2018, Gordon & Riley, 1992, Schreiber *et al*., 2017, Gibreel *et al*., 2012, Sintsova *et al*., 2019). For instance, studies focused on identifying characteristics of uropathogenic *E. coli* isolates underlying their propensity for causing infection in comparison to fecal isolates or asymptomatic colonizers have consistently revealed a role for metabolism and rapid growth in urine (Forsyth *et al*., 2018, Schreiber *et al*., 2017, Gibreel *et al*., 2012, Sintsova *et al*., 2019). In addition, genome-wide screens in other common uropathogens have similarly pointed to the importance of metabolic networks for adaptation to growth in urine and fitness within the urinary tract (Armbruster *et al*., 2017a, Johnson *et al*., 2020, Tielen *et al*., 2013, Paudel *et al*., 2021, Colomer-Winter *et al*., 2019). Thus, investigating the metabolic pathways that facilitate rapid growth in urine may provide new insights for combating UTIs.

Normal human urine represents a high-salt, low-iron, nutrient-limited medium in which the most abundant catabolites are generally urea, citrate, amino acids, and small peptides (Bouatra *et al*., 2013, Brooks & Keevil, 1997, Reitzer & Zimmern, 2019). A common finding from the genome-wide screens in uropathogenic bacteria is that growth in urine induces a shift from sugar catabolism to amino acid import and metabolism, indicating that the ability to use amino acids as carbon and nitrogen sources is important for urinary tract colonization. The most abundant amino acids in urine are typically glycine, histidine, alanine, serine, glutamine, threonine, asparagine, and lysine (Dunstan *et al*., 2017, Armbruster *et al*., 2013, 2021). Urine is also unique from other bodily fluids as it contains a high concentration of D-serine in addition to the L- amino acid enantiomers, with the ratio of D- to L-serine being approximately 1:1 (Miyoshi *et al*., 2009, Bruckner & Schieber, 2001, Brauer *et al*., 2019). It has therefore been hypothesized that the ability to import and catabolize D-serine may provide a substantial benefit during urinary tract colonization. Not all uropathogens are capable of catabolizing D-serine, but genes for D-serine import and catabolism are generally more prevalent in uropathogenic isolates of several species (Cosloy & McFall, 1973, Nørregaard-Madsen *et al*., 1995, Korte-Berwanger *et al*., 2013, Sakinç *et al*., 2009).

*P. mirabilis* is one of the most common uropathogens, particularly in older adults and individuals with urinary catheters (Gaston *et al*., 2021, Armbruster *et al*., 2021, Armbruster *et al*., 2017b). This organism is of particular concern as it is highly persistent within the urinary tract and notoriously difficult to treat due to a combination of intrinsic drug resistances, acquisition of extended-spectrum β-lactamase and carbapenemase, and its ability to form recalcitrant crystalline biofilms (Armbruster *et al*., 2018). It is also one of the most common causes of infection-induced urinary stones and a leading cause of bacteremia and mortality in catheterized individuals (Griffith *et al*., 1976, Li *et al*., 2002, Foxman & Brown, 2003, Kim *et al*., 2003, Watanakunakorn & Perni, 1994). Given the serious complications that can arise from *P. mirabilis* colonization and infection and increasing antimicrobial resistance, we and others have performed genome-wide screens to identify key fitness factors of *P. mirabilis* that facilitate colonization and persistence within the urinary tract (Burall *et al*., 2004, Himpsl *et al*., 2008, Zhao *et al*., 1999, Pearson *et al*., 2011, Armbruster *et al*., 2017a, Armbruster *et al*., 2019). Similar to studies of uropathogenic *E. coli*, a common theme in these studies has been the identification of numerous metabolic pathways that appear to provide niche-specific fitness within the urinary tract, including genes for the import and utilization of D- and L-serine along with other amino acids.

We previously determined that *P. mirabilis* strain HI4320 can utilize D-serine as a sole source of carbon and nitrogen, and that D-serine utilization contributes to colonization and fitness within the catheterized urinary tract (Brauer *et al*., 2019). However, the contribution of L-serine to colonization and fitness had yet to be explored. The goal of this study was to determine the full hierarchy of amino acid utilization by *P. mirabilis* during growth in urine and to examine the relative importance of D- vs L-serine catabolism for critical steps in CAUTI development and progression, including: 1) overall fitness compared to wild-type *P. mirabilis,* 2) swimming and swarming motility for migration along the catheter surface and ascension of the ureters, 3) growth in urine, 4) biofilm formation for development of persistent bacterial communities, 5) infection of the bladder and kidneys, and 6) dissemination to the bloodstream.

## Results

### *P. mirabilis* preferentially depletes serine over other abundant amino acids during growth in urine

High-performance liquid chromatography (HPLC) was used to determine amino acid concentrations in pooled normal human urine (Table 1). The most abundant amino acids overall were glycine, alanine, and arginine, followed by histidine and methionine. The concentration of most amino acids remained stable in uninoculated human urine during incubation at 37°C for 12 hours (Fig 1A), while supernatants from urine inoculated with *P. mirabilis* revealed depletion below the limit of detection for seven amino acids (threonine, glutamine, serine, tyrosine, tryptophan, phenylalanine, and isoleucine) and a 50% reduction of five additional amino acids (histidine, glutamic acid, arginine, leucine, and asparagine) (Fig 1B). One notable exception was methionine, which *P. mirabilis* appears to produce and secrete over time, resulting in a final concentration roughly 1000% of the starting concentration (Fig 1B).

**Figure 1.**
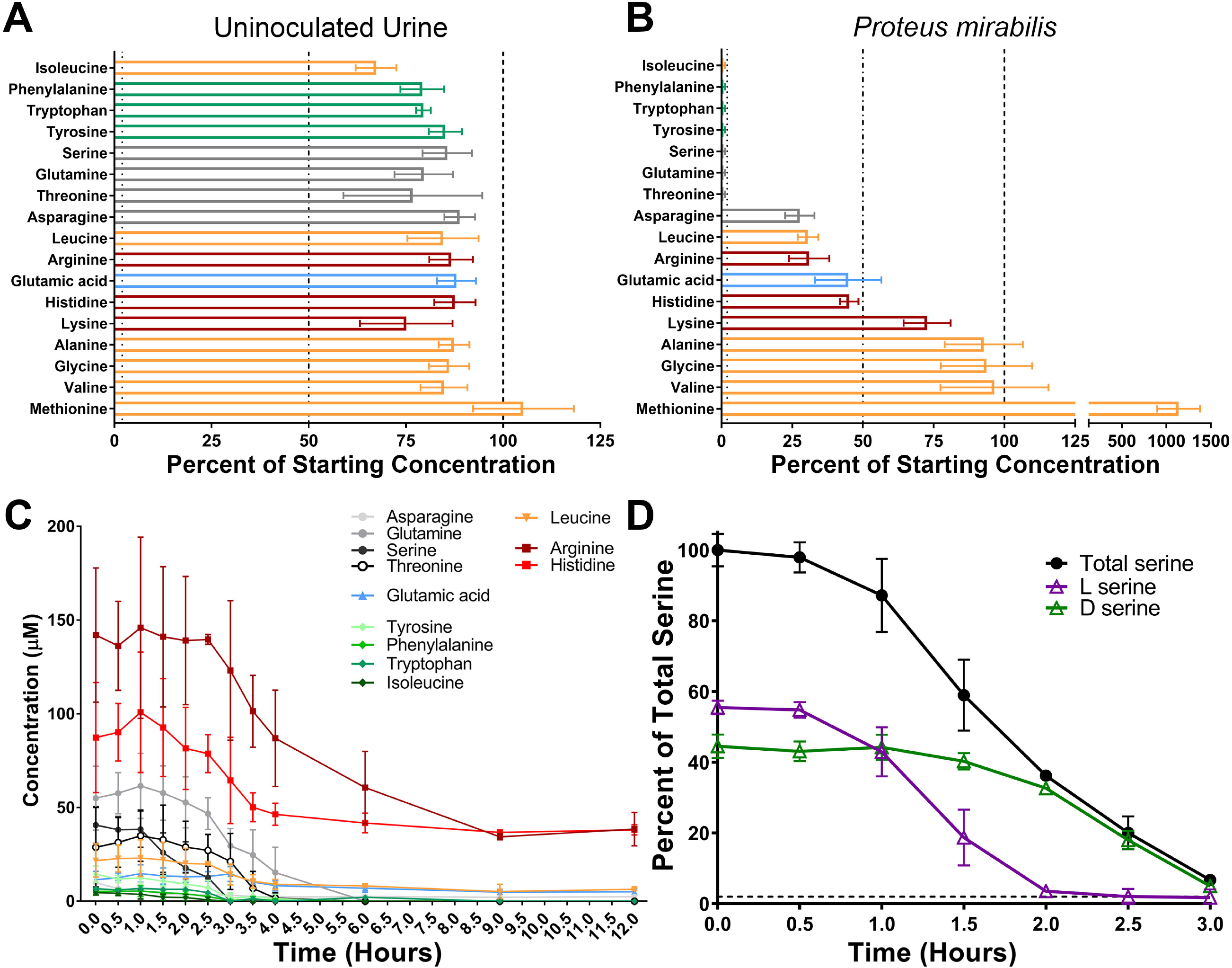
*P. mirabilis* preferentially depletes L-serine and D-serine before other amino acids during growth in human urine. Filter-sterilized pooled human urine from at least 20 female donors was inoculated with wild-type *P. mirabilis* HI4320 and incubated at 37° with aeration for 12 hours, with aliquots removed every half hour for quantification of free amino acids in cell-free supernatants by HPLC. (A and B) Data indicate the percentages of 17 amino acids after 12 hours of incubation for uninoculated urine (A) and urine inoculated with *P. mirabilis* (B). Categories of amino acids are color-coded as follows: polar uncharged (gray); negatively charged (blue); aromatic (green); non-polar aliphatic (orange); positively charged (red). Error bars represent mean ± standard deviation (SD) from 3 independent experiments, and dashed lines indicate 100%, 50%, and 1%. (C) Concentrations of all 12 amino acids with that were depleted at least 50% in panel B are displayed over time. Error bars represent mean ± standard deviation (SD) from 3 independent experiments. (D) Replicate supernatants from the experiment in panel C were used to determine the relative amount of each serine enantiomer over time. Data are expressed as percentage of total serine (black lines). Error bars represent mean ± SD for 3 independent experiments with 3 technical replicates each.

**Table 1:** Concentration of amino acids in human urine.

|  | **Concentration**<br>**(μM)<sup>a</sup>** | **Standard**<br>**Deviation** |
| --- | --- | --- |
| **Glycine** | 184.8 | 54.3 |
| **Alanine** | 139.0 | 37.8 |
| **Arginine** | 133.6 | 37.9 |
| **Histidine** | 85.5 | 27.1 |
| **Methionine** | 84.7 | 29.8 |
| **Glutamine** | 53.9 | 15.9 |
| **Serine** | 39.1 | 10.6 |
| **Valine** | 39.2 | 10.7 |
| **Lysine** | 34.9 | 6.5 |
| **Threonine** | 22.5 | 9.1 |
| **Leucine** | 20.0 | 7.5 |
| **Tyrosine** | 13.3 | 3.6 |
| **Glutamic acid** | 11.4 | 3.9 |
| **Asparagine** | 11.4 | 3.7 |
| **Tryptophan** | 6.0 | 2.1 |
| **Phenylalanine** | 4.8 | 1.7 |
| **Isoleucine** | 3.9 | 2.5 |
| **Aspartic acid** | ND |  |
| **Cysteine** | ND |  |
| **Proline** | ND |  |

<sup>a</sup>Mean concentration detected by HPLC from 3 biological replicates with 3 technical replicates each. The limit of detection was ~1 μM. ND=not detected or below the limit of detection.

In order to determine the timing of depletion, we monitored the concentration of each amino acid every half hour for 4 hours and again at 6, 9, and 12 hours. Serine and isoleucine were depleted first (∼3 hours), followed by threonine and tyrosine (∼4 hours), then glutamine, tryptophan, and phenylalanine (∼6 hours) (Fig 1C). Notably, threonine and glutamine levels only began to decrease after serine had been fully depleted. To determine if *P. mirabilis* preferentially depletes one serine enantiomer over the other, we next assessed the timing of depletion of L- vs D-serine. Strikingly, D-serine levels only began to decrease after L-serine had been depleted (Fig 1D). Taken together, these data indicate that *P. mirabilis* exhibits a clear hierarchy of amino acid import during growth in urine and a preferential utilization of L- vs D-serine.

### *P. mirabilis* strain HI4320 can utilize L-serine as a sole carbon or nitrogen source through the combined action of two L-serine deaminases

*E. coli* genomes typically contain three enzymes capable of producing pyruvate and ammonia from L-serine (*sdaA, sdaB,* and *tdcG*), of which SdaA and SdaB exhibit ∼77% amino acid identity to each other and 73-74% amino acid identity to TdcG (Heßlinger *et al*., 1998, Zhang & Newman, 2008). In contrast, a basic local alignment search of the genome of *P. mirabilis* strain HI4320 identified only two such enzymes; *sdaA* (PMI1607) and *sdaB* (PMI0671). We therefore disrupted each gene alone and created a double mutant to assess their relative contributions to L-serine catabolism in *P. mirabilis*.

All three mutants exhibited comparable growth to wild-type *P. mirabilis* in minimal medium containing glycerol as the carbon source and ammonium sulfate as the nitrogen source (Fig 2A). When *P. mirabilis* was forced to use L-serine as either the sole carbon (Fig 2B) or nitrogen source (Fig 2C), growth was reduced by loss of *sdaA* and completely inhibited by loss of both *sdaA* and *sdaB,* but unaffected by the loss of *sdaB* alone. The growth defect of the *sdaAB* double mutant could be largely restored via complementation with either *sdaA* or *sdaB* on a plasmid (Fig 2D). These data demonstrate that *P. mirabilis* can utilize L-serine as either a sole carbon or nitrogen source, that *sdaA* and *sdaB* are the only functional L-serine deaminases in strain HI4320, and that *sdaA* may be favored in wild-type *P. mirabilis* under these experimental conditions.

**Figure 2.**
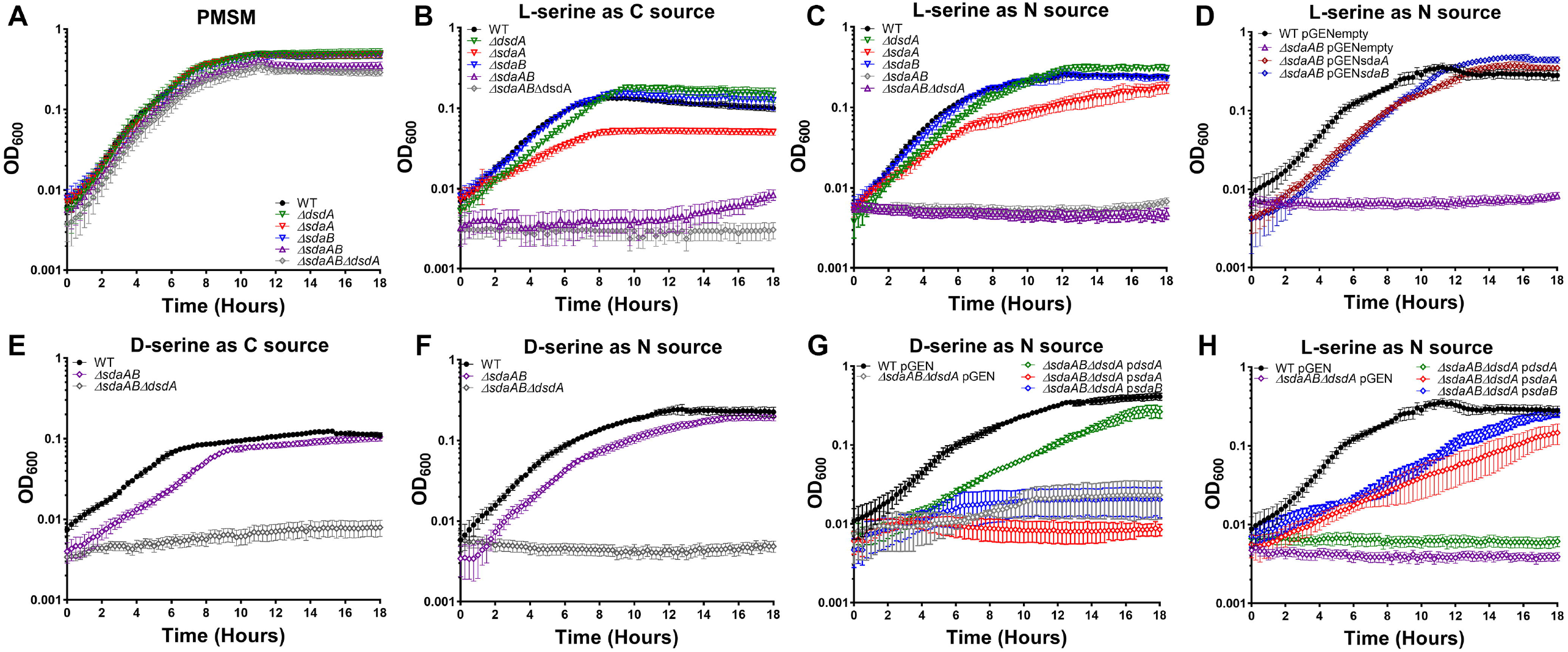
Either L-serine deaminase is sufficient for utilization of L-serine as a sole carbon or nitrogen source, and *sdaA, sdaB,* and *dsdA* encode the only functional serine catabolic genes in *P. mirabilis* HI4320. Wild-type *P. mirabilis* and indicated serine catabolism mutants were cultured in minimal medium (PMSM) with the following sole carbon/nitrogen sources: (A) glycerol/ammonium sulfate; (B) L-serine/ammonium sulfate; (C, D, and H) glycerol/L-serine; (E) D-serine/ammonium sulfate; (F and G) glycerol/D-serine. Cultures were incubated at 37°C with aeration in 96-well plates, and growth was assessed by measurement of OD_600_ at 15-minute intervals for 18 hours. Error bars represent mean ± SD from at least 6 technical replicates, and graphs are representative of at least 3 independent experiments. Where indicated, mutants were complemented by electroporation with a pGEN plasmid empty vector (pGENempty) or harboring the indicated gene (e.g. p*sdaA*).

We previously demonstrated that *P. mirabilis* can use D-serine as a sole carbon or nitrogen source due to D-serine dehydratase (*dsdA*) (Brauer *et al*., 2019). To verify that *sdaA, sdaB,* and *dsdA* are the only genes responsible for catabolism of either serine enantiomer, a triple mutant was constructed and assessed for growth with either D- or L-serine as the sole carbon or nitrogen source. The triple mutant grew similarly to the *sdaAB* double mutant and wild-type *P. mirabilis* in minimal medium containing glycerol as the carbon source and ammonium sulfate as the nitrogen source (Fig 2A), but was unable to grow when either D- or L-serine were provided as the sole carbon or nitrogen source (Fig 2 B, C, E, and F). Growth with D-serine as the nitrogen source was partially restored by providing *dsdA* on a plasmid (Fig 2G), and growth with L-serine as the nitrogen source was partially restored by providing either *sdaA* or *sdaB* on a plasmid (Fig 2H). Taken together, these data indicate that loss of serine catabolism does not impair growth of *P. mirabilis* when glycerol and ammonium are supplied as the carbon and nitrogen sources, and that *dsdA, sdaA,* and *sdaB* encode the only serine catabolism enzymes in strain HI4320.

We next sought to identify and experimentally confirm the serine importers encoded by *P. mirabilis* strain HI4320. This isolate encodes a single D-serine transporter (*dsdX*, PMI0186) which we previously determined to be the only functional D-serine importer (Brauer *et al*., 2019), as well as two putative L-serine/threonine importers; *sdaC* (PMI0672) and *sstT* (PMI3701). Both genes were previously predicted to be serine transporters and tested for their contribution to self-recognition during *P. mirabilis* swarming (Chittor & Gibbs, 2021), although individual deletion mutants were not generated to verify their contribution to serine or threonine import. We therefore disrupted each gene alone and created a double mutant to assess their relative contributions to amino acid import in *P. mirabilis.* All three mutants exhibited comparable growth to wild-type *P. mirabilis* in minimal medium containing glycerol as the carbon source and ammonium sulfate as the nitrogen source (Fig 3A). When L-serine was supplied as the only nitrogen source, a noticeable defect was observed for the *sstT* mutant and the *sstTsdaC* but all three importer mutants remained capable of at least modest growth (Fig 3B), indicating that *P. mirabilis* HI4320 imports serine through at least one additional transport system.

**Figure 3.**
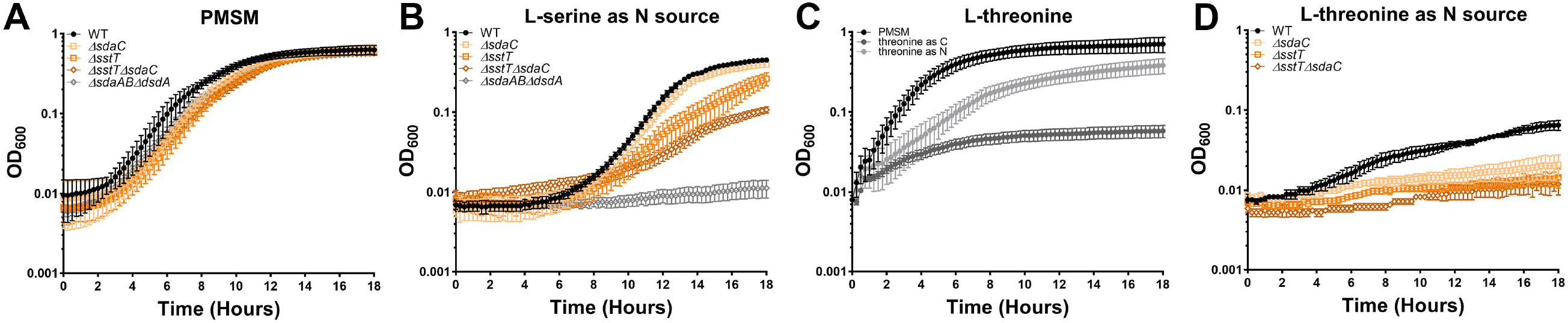
*P. mirabilis* encodes multiple transporters that contribute to both serine and threonine import. Wild-type *P. mirabilis* and indicated serine/threonine import mutants were cultured in standard PMSM or formulations containing either L-serine or L-threonine as the sole carbon or nitrogen source. (A) Growth of wild-type *P. mirabilis,* serine/threonine import single mutants, and *sstTsdaC* double mutant in standard PMSM. (B) Growth of wild-type *P. mirabilis,* serine/threonine import single mutants, and *sstTsdaC* double mutant in PMSM containing L-serine as the sole nitrogen source. The *sdaABdsdA* mutant was included as a no-growth control. (C) Growth of wild-type *P. mirabilis* was compared across standard PMSM and PMSM containing L-threonine as the sole carbon or nitrogen source. (D) Growth of wild-type *P. mirabilis,* serine/threonine import single mutants, and *sstTsdaC* double mutant in PMSM containing L-threonine as the sole nitrogen source. All cultures were incubated at 37°C with aeration in 96-well plates, and growth was assessed by measurement of OD_600_ at 15-minute intervals for 18 hours. Error bars represent mean ± SD from at least 6 technical replicates, and graphs are representative of at least 3 independent experiments.

Since *sdaC* and *sstT* are also annotated as potential threonine importers, we next sought to determine their contribution to L-threonine import. Wild-type *P. mirabilis* exhibited modest growth when supplied with L-threonine as a sole nitrogen source and failed to survive when forced to use it as the sole carbon source (Fig 3C). Disrupting either *sdaC* or *sstT* substantially impaired growth on L-threonine as the sole nitrogen source, and the double mutant completely failed to grow (Fig 3D). These data indicate that *sdaC* and *sstT* are capable of importing both L-serine and L-threonine, but additional transporters likely contribute to serine import in *P. mirabilis* strain HI4320. Due to this complication, the rest of this body of work focuses on serine catabolism mutants to ensure specificity.

### Disrupting serine catabolism decreases fitness of *P. mirabilis* during growth in LB

Prior studies in *E. coli* have highlighted important potential caveats to studying serine deaminase mutants. For instance, loss of L-serine catabolism in *E. coli* causes intracellular serine accumulation (Anfora *et al*., 2007), and the resulting excess L-serine can interfere with optimal growth when rich carbon sources like glucose or carbohydrates are present (Kriner & Subramaniam, 2020, Zhang *et al*., 2010). This has been hypothesized to occur through several potential mechanisms, including inhibition of isoleucine, threonine, and aromatic amino acid biosynthesis (Hama *et al*., 1991, Tazuya-Murayama *et al*., 2006), starvation for single-carbon units due to inhibition of serine hydroxymethyl transferase (GlyA) and glycine cleavage (Zhang & Newman, 2008), and disruption of cell wall integrity due to competition between L-serine and L-alanine for MurC during generation of peptidoglycan (Parveen & Reddy, 2017). Serine levels and the serine transporter SdaC have also been proposed to play a role in maintaining amino acid homeostasis during metabolic shifts in *E. coli* (Kriner & Subramaniam, 2020). In *P. mirabilis,* prior work in strain BB2000 demonstrated that disruption of *sdaAB* lead to the accumulation of intracellular serine in swarmer cells (Chittor & Gibbs, 2021). For strain HI4320, we previously demonstrated that loss of D-serine catabolism via disruption of *dsdA* causes accumulation of intracellular D-serine and subsequent disruption of L-serine and pantothenate biosynthesis, which can be remedied by either the addition of excess L-serine or pantothenic acid (Brauer *et al*., 2019). We therefore sought to determine the impact of disrupting D- and L-serine catabolism on *P. mirabilis* growth rate in Lysogeny Broth (LB, Luria recipe) as a representative rich medium containing carbohydrates, peptides, and free amino acids.

All serine utilization mutants grew similarly to wild-type *P. mirabilis* in LB, exhibiting at most a modest decrease in cell density during exponential phase (Fig 4A). However, clear differences were uncovered by directly competing the mutants against wild-type *P. mirabilis*. While no competition was observed between wild-type *P. mirabilis* and the D-serine catabolism mutant, loss of L-serine catabolism resulted in a significant fitness defect as early as 4 hours post-inoculation and the combined loss of both D- and L-serine catabolism resulted in an even greater fitness defect that was apparent as early as 1 hour post-inoculation (Fig 4B). Thus, disrupting serine catabolism perturbs *P. mirabilis* fitness in rich medium as previously reported in *E. coli,* albeit to a lesser extent.

**Figure 4.**
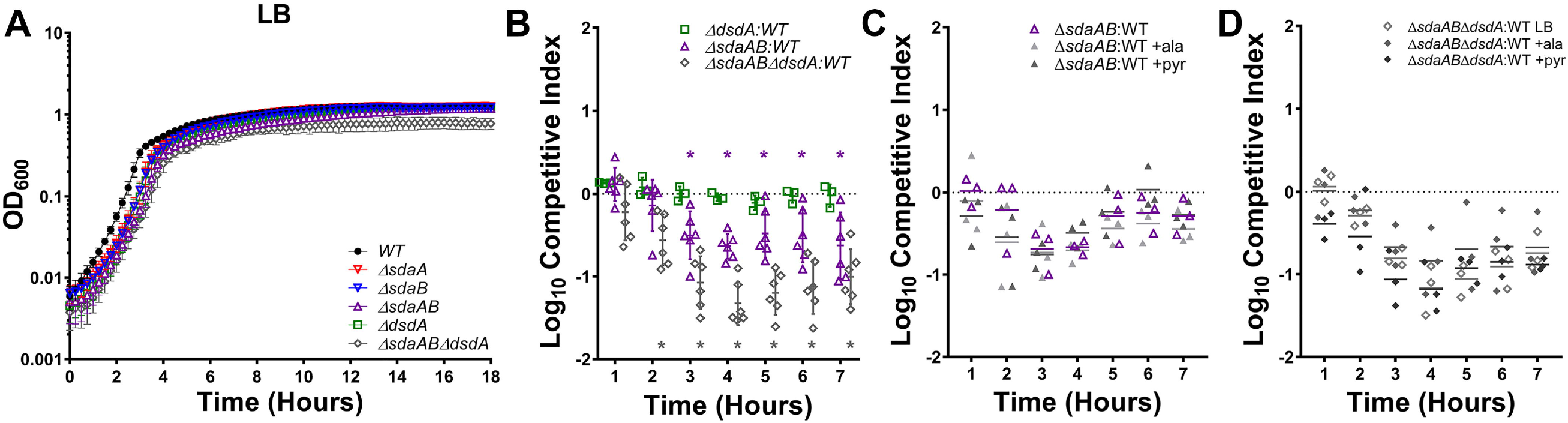
L-serine catabolism contributes to *P. mirabilis* fitness in LB broth. (A) Wild-type *P. mirabilis* and indicated serine catabolism mutants were cultured in LB broth at 37°C with aeration in 96-well plates, and growth was assessed by measurement of OD_600_ at 15-minute intervals for 18 hours. Error bars represent mean ± SD from at least 6 technical replicates, and graph is representative of at least 3 independent experiments. (B-D) LB broth was inoculated with a 50:50 mixture of the indicated serine catabolism mutant and wild-type *P. mirabilis,* and fitness was assessed by calculating a competitive index based on the CFUs of each strain at hourly time points. Where indicated, LB broth was supplemented with 10 mM alanine or pyruvate. Each data point indicates the log_10_ competitive index for the indicate mutant:WT inoculum from a single culture. Error bars represent the mean ± SD from at least 3 independent experiments, and the dashed line indicates Log_10_ CI=0 (the expected value if the ratio of mutant:WT is 1:1). \**p*<0.05 by Wilcoxon signed-rank test, with specific co-challenges indicated by color-coding.

We hypothesized that the fitness defects resulting from loss of serine catabolism could be due to one of three main factors: 1) loss of energy production from serine due to lack of catabolism to pyruvate, 2) perturbation of cell wall integrity due to competition between serine and alanine during peptidoglycan synthesis, or 3) general perturbation of amino acid homeostasis due to intracellular serine accumulation. To determine which factors may be responsible for the growth defects of the serine mutant, we supplemented LB with 10 mM pyruvate to correct any issues related to energy production or with 10 mM alanine to address issues pertaining to amino acid homeostasis and cell wall integrity. Supplementation with either pyruvate or alanine failed to rescue the fitness defect of either the *sdaAB* double mutant (Fig 4C) or the *sdABdsdA* triple mutant (Fig 4D), indicating that the fitness defect in LB broth that results from loss of L-serine catabolism is not due to competition between serine and alanine during peptidoglycan synthesis or loss of energy production from serine.

### Complete loss of serine catabolism disrupts *P. mirabilis* membrane integrity during growth on media with a rich carbon source

To further explore the source of decreased fitness in the serine catabolism mutants, we next assessed overall cell wall and membrane integrity by subjecting wild-type *P. mirabilis* and each of the mutants to discs soaked in the following agents: an antimicrobial agent that targets DNA replication to which increased susceptibility would suggest a change in cell wall or membrane permeability (ciprofloxacin), antimicrobials that target peptidoglycan biosynthesis (imipenem and ceftazidime), and detergents (sodium deoxycholate [DOC] and sodium dodecyl sulfate [SDS]). Disrupting either D- or L-serine catabolism alone had no impact on sensitivity to any of the agents, indicating that any potential serine accumulation that occurs in these mutants during growth on low-salt LB agar is insufficient to perturb cell integrity (Fig 5A-E). However, the combined loss of both serine catabolism pathways increased susceptibility to all five agents (Fig 5A-E). Plasmid-based complementation experiments were not conducted due to potential issues with maintaining plasmid selection in *P. mirabilis* during exposure to antimicrobial agents and detergents, but supplementation with either 10 mM pyruvate or alanine failed to reverse sensitivity of the triple mutant (Fig 5F-I). Since excess alanine should counteract any potential cell wall biosynthesis defects resulting from intracellular accumulation of serine, these data suggest that the complete loss of serine catabolism may perturb cell integrity through an alternative mechanism than what has been previously described in *E. coli*.

**Figure 5.**
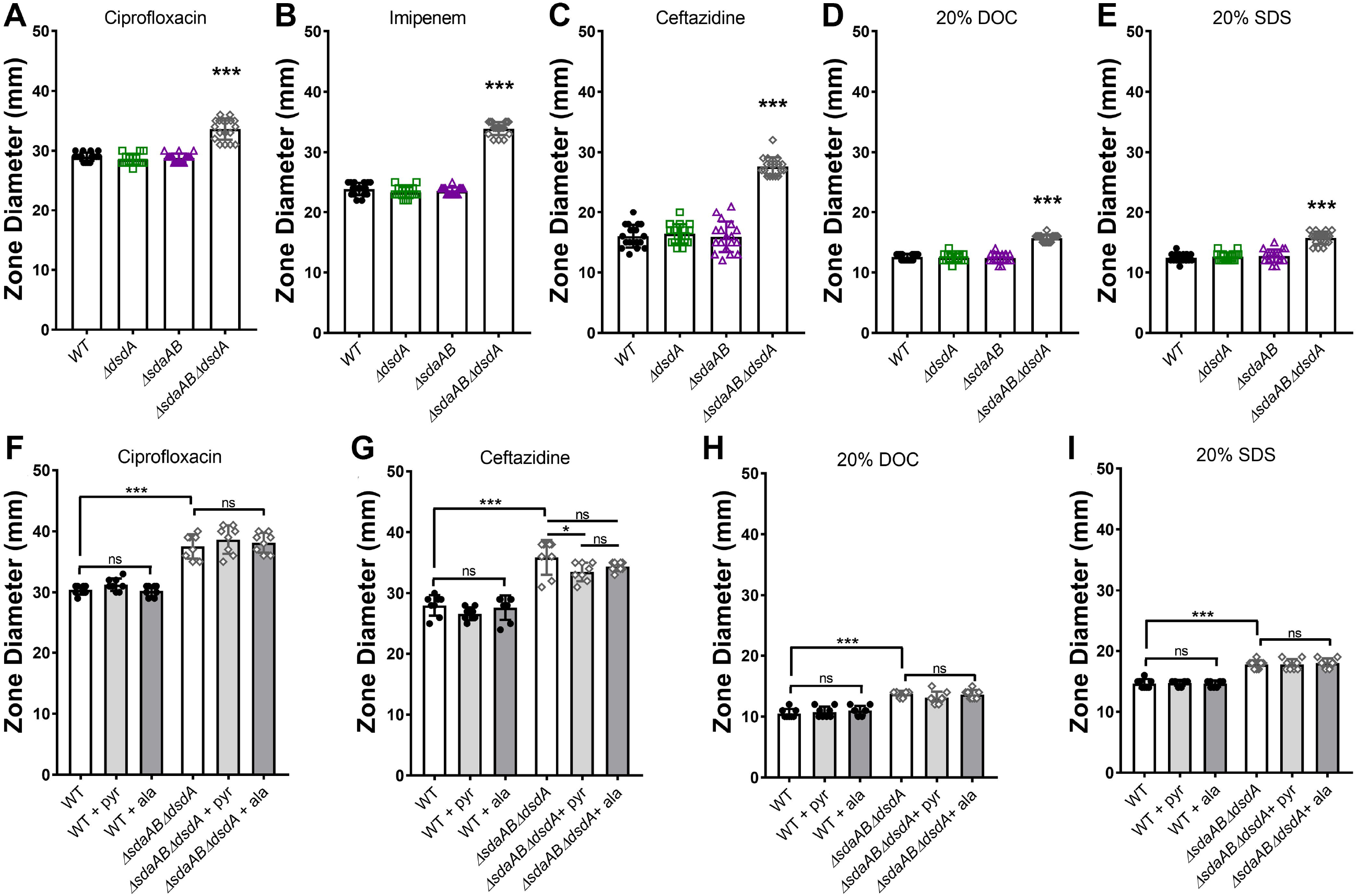
Complete disruption of serine catabolism increases sensitivity to antimicrobial agents and detergents. Wild-type *P. mirabilis* and indicated mutants were swabbed across the surface of LB agar plates to generate bacterial lawns, and discs impregnated with the following agents were placed on the plates to quantify zones of inhibition: (A and F) 5µg ciprofloxacin; (B) 10µg imipenem; (C and G) 30µg ceftazidime; (D and H) 20% sodium dodecyl sulfate; (E and I) 20% sodium deoxycholate. Plates were incubated at 37°C for 18 hours, after which the diameter of the zone of inhibition was measured. Where indicated, LB broth was supplemented with 10 mM alanine or pyruvate. Error bars represent mean ± SD from at least 3 independent experiments with 3 technical replicates each. \*\*\**p*<0.001 by one-way ANOVA with Dunnett’s test for multiple comparisons. ns=not significant.

1-*N*-phenylnaphthylamine (NPN) is a hydrophobic dye that fluoresces in phospholipid-rich environments, such as the inner leaflet of the outer membrane in Gram-negative bacteria, thereby permitting assessment of membrane permeability (Helander & Mattila-Sandholm, 2000). To determine if loss of serine catabolism specifically alters membrane permeability, NPN fluorescence was measured in the *dsdA, sdaAB,* and *sdaABdsdA* mutants for comparison to wild-type *P. mirabilis* with or without the membrane permeabilizer ethylenediaminetetraacetic acid (EDTA, Fig 6A). Exposure to 62.5 mM EDTA slightly increased *P. mirabilis* membrane permeability, resulting in an increase in NPN uptake as measured by fluorescence. Loss of D-serine catabolism alone did not impact membrane permeability, but loss of L-serine catabolism resulted in a greater degree of NPN uptake than treatment with EDTA and the combined loss of both D- and L-serine catabolism further increased permeability. Thus, disrupting serine catabolism alters *P. mirabilis* membrane permeability.

**Figure 6.**
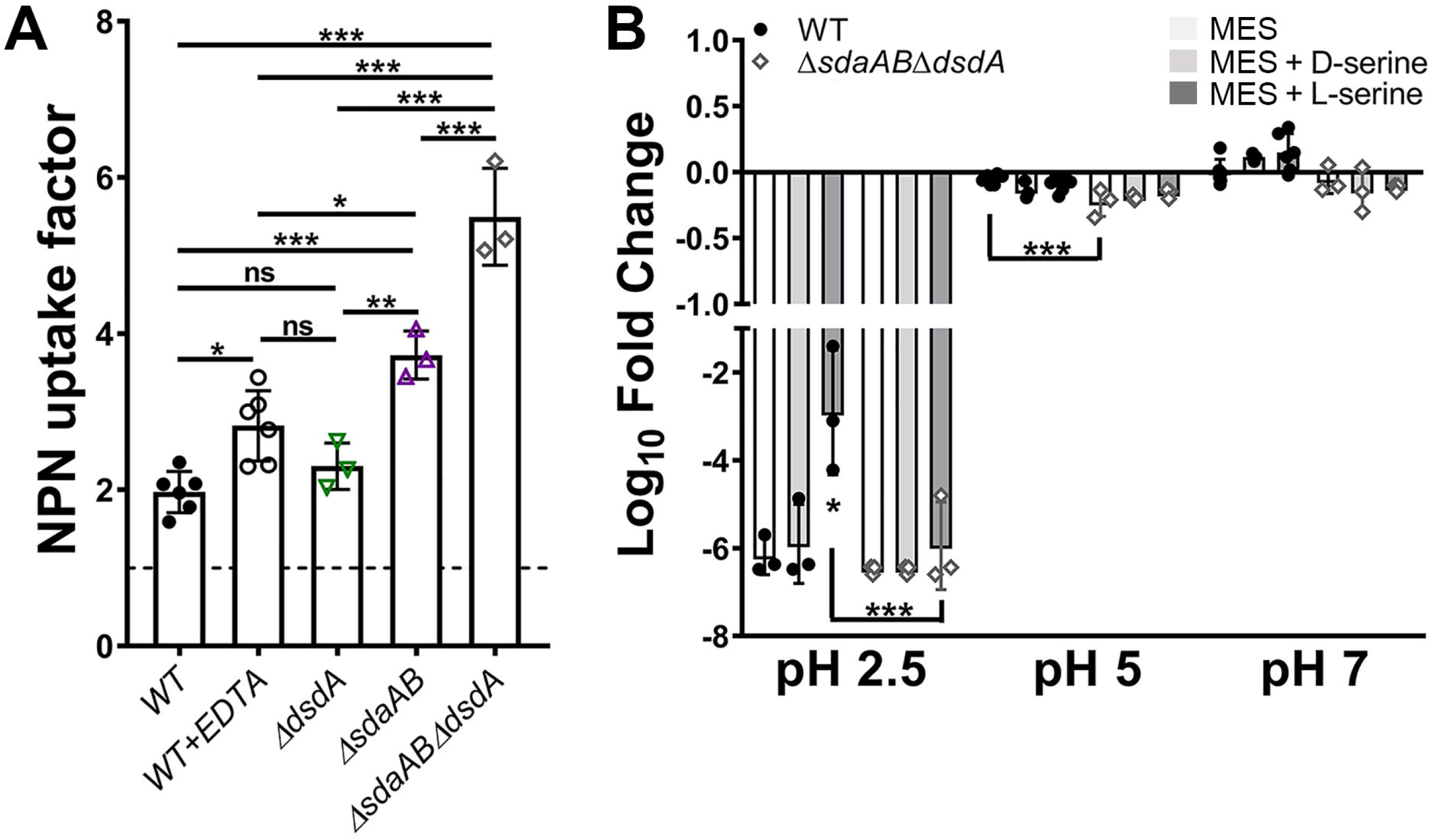
Disruption of L-serine catabolism increases membrane permeability. (A) Mid-log cultures of wild-type *P. mirabilis* and indicated mutants were adjusted to the same starting density in HEPES buffer and incubated with 1-*N*-phenylnaphthylamine (NPN), and an NPN uptake factor was calculated based on the fluorescence from background-subtracted samples and controls as described in the Methods section. Treatment with EDTA was included as a positive control for increased membrane permeability. Error bars represent mean ± SD from at least 3 independent experiments. \**p*<0.05, \*\**p*<0.01, \*\*\**p*<0.001 by one-way ANOVA with Dunnett’s test for multiple comparison. ns=not significant. (B) Mid-log cultures of wild-type *P. mirabilis* and the *sdaABdsdA* triple mutant were adjusted to the same starting density in MES buffer at pH 2.5, 5, or 7 and incubated at 37°C for 1 hour prior to plating for determination of CFUs. Where indicated, MES was supplemented with 10 mM D-serine (light gray bars) or L-serine (dark gray bars). Error bars represent mean ± SD from 3 independent experiments. \**p*<0.05, \*\*\**p*<0.001 by two-way ANOVA with Tukey’s test for multiple comparisons.

Changes in membrane permeability can also alter membrane potential and proton motive force, which are critical components of acid tolerance and motility in bacteria. We previously demonstrated that catabolism of another amino acid, arginine, contributes to maintenance of membrane potential in *P. mirabilis* due to the consumption of intracellular protons (H^+^), and that acid tolerance can be utilized as surrogate for assessing changes in membrane potential in *P. mirabilis* strain HI4320 (Armbruster *et al*., 2014). While the serine deaminase reaction does not directly consume a proton, pyruvate is involved in numerous proton-consuming reactions that may ultimately influence membrane potential. Wild-type *P. mirabilis* and the *sdaABdsdA* mutant were therefore assessed for survival after a 60-minute incubation at pH 2.5, 5, and 7 to determine if complete loss of serine catabolism impacts membrane potential (Fig 6B). Consistent with our prior observations, the viability of wild-type *P. mirabilis* was reduced almost to the limit of detection during incubation at pH 2.5 but largely unaffected at pH 5 or 7. Supplementation with D-serine had no impact on *P. mirabilis* viability while L-serine increased viability during incubation at pH 2.5, suggesting that L-serine catabolism may contribute to maintenance of membrane potential under these conditions. The Δ*sdaAB*Δ*dsdA* mutant exhibited similar viability trends as wild-type *P. mirabilis* at each pH, with the exception of a modest though statistically significant decrease in viability at pH 5. However, viability of the Δ*sdaAB*Δ*dsdA* mutant was not augmented by supplementation with L-serine, providing additional evidence that serine catabolism may contribute to maintenance of membrane potential in *P. mirabilis* strain HI4320.

### Serine catabolism contributes to flagellar-mediated motility in *P. mirabilis*

*P. mirabilis* is capable of robust swimming motility due to its peritrichous flagella as well as flagellum-mediated swarming motility, both of which contribute to pathogenesis by facilitating ascension of the urethra and ureters (Armbruster *et al*., 2018). We previously demonstrated that D-serine import and catabolism are not required for *P. mirabilis* motility (Brauer *et al*., 2019), but the L-serine transporter SdaC was recently linked to swarm expansion (Chittor & Gibbs, 2021). In *P. mirabilis* strain BB2000, SdaC was important for proper coordination of swarming while the combined loss of *sdaA* and *sdaB* abrogated swarming, which the authors hypothesized was due to intracellular accumulation of L-serine in swarmer cells of the double mutant. However, considering that flagella-mediated motility is dependent upon membrane potential and proton motive force, and these are both perturbed in serine catabolism mutants, we sought to further explore the relative contribution of D- and L-serine catabolism to swarming and swimming motility in *P. mirabilis* strain HI4320.

Disruption of D-serine utilization alone had no impact on swarming motility (Fig 7A), confirming our previous results (Brauer *et al*., 2019). Disruption of L-serine utilization in *P. mirabilis* strain HI4320 resulted in only a modest decrease in the diameter of each individual swarm ring (Fig 7A), which may represent a strain-dependent difference for HI4320 compared to BB2000. However, the combined loss of both D- and L-serine catabolism further impaired swarming, suggesting that either serine catabolism is important for swarm migration or that accumulation of intracellular serine interferes with motility. Swarming defects were substantially more pronounced in strains carrying pGEN under ampicillin selection, but the defects of the L-serine utilization mutant could be fully complemented by providing either *sdaA* or *sdaB* on a plasmid, and migration of the triple mutant was largely restored by providing either *dsdA, sdaA,* or *sdaB* on a plasmid (Fig 7B). Supplementation with alanine also restored swarming to roughly wild-type levels for both Δ*sdaAB* and Δ*sdaAB*Δ*dsdA* while pyruvate had no impact (Fig 7C), suggesting that the motility defects are largely due to general perturbation of amino acid homeostasis and competition between serine and alanine during cell wall biosynthesis.

**Figure 7.**
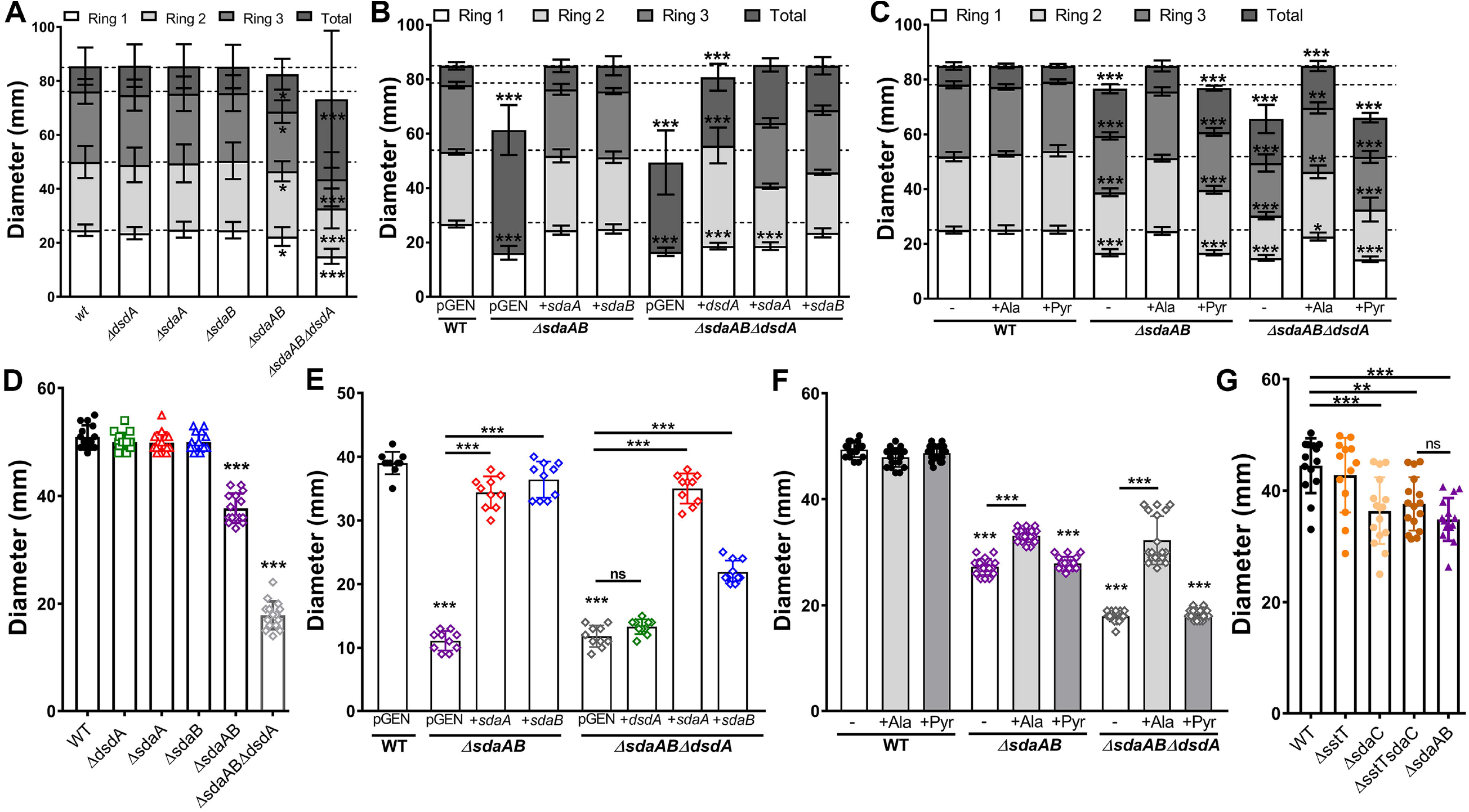
Disruption of L-serine catabolism perturbs swarming and swimming motility. Wild-type *P. mirabilis* and indicated mutants were inoculated onto the surface of swarm agar plates (A-C) or stab inoculated into motility agar (D-G) as follows: (A, D, and G) standard media formulations; (B and E) media containing 100 µg/mL ampicillin to maintain plasmid selection; (C and F) media supplemented with 10 mM alanine or pyruvate. Motility diameter was measured in millimeters after 16 hours of incubation at 37°C (A-C) or 30°C (D-F). Error bars represent mean ± SD for at least three independent experiments with at least 3 replicates each. \**p*<0.05, \*\**p*<0.01, \*\*\**p*<0.001 by two-way ANOVA with Tukey’s test for multiple comparisons (A-C) or by one-way ANOVA with Dunnett’s test for multiple comparison (D-G).

Since swarming represents a coordinated multicellular behavior in *P. mirabilis,* we next assessed swimming motility to determine if the swarming defects were representative of a broad motility defect or if they specifically pertain to multicellular migration on a solid surface. Disruption of D-serine utilization alone had no impact on motility (Fig 7D), again confirming our previous results (Brauer *et al*., 2019). Disruption of either *sdaA* or *sdaB* alone had no impact on motility, but complete loss L-serine utilization in the *sdaAB* double mutant resulted in a substantial motility defect and the combined loss of both D- and L-serine utilization further impaired motility (Fig 7D). Motility defects were again more pronounced in strains carrying pGEN under ampicillin selection, but the swimming defect of the L-serine utilization mutant was fully complemented by providing either *sdaA* or *sdaB* on a plasmid (Fig 7E). In contrast to the swarming motility complementation experiments, the motility defect of the *sdaABdsdA* mutant could only be restored by providing *sdaA* on a plasmid, while *sdaB* provided only partial complementation and *dsdA* had no effect (Fig 7E). Similarly, supplementation with alanine only partially restored motility and pyruvate was ineffective (Fig 7F). To further delineate the underlying cause of the swimming motility defects, we next assessed motility in the in L-serine import mutants *sdaC, sstT,* and a double mutant (Fig 7G). The *sdaC* and *sstTsdaC* double mutant both exhibited swimming motility defects that were almost identical to the *sdaAB* double mutant. Collectively, these data indicate that serine import and catabolism directly contribute to swimming motility due to a role in membrane permeability and proton motive force, and that any serine accumulation in the catabolism mutants only exerts a modest effect due to competition between serine and alanine during cell wall biosynthesis.

### Catabolism of either D- or L-serine is sufficient for fitness during growth in human urine but L-serine catabolism is favored

Consistent with studies in *E. coli,* the most pronounced defects of the serine mutants occurred during growth in LB rather than nutrient-limited minimal medium. Considering that urine is a nutrient-limited medium lacking carbohydrates and rich carbon sources and that *P. mirabilis* preferentially depletes L-serine followed by D-serine before other amino acids during growth in urine (Fig 1A), we next sought to determine the relative contribution of D- and L-serine degradation to *P. mirabilis* growth and fitness in human urine (Fig 8). The *dsdA* mutant grew similarly to wild-type *P. mirabilis* in human urine (Fig 8A), confirming our prior results (Brauer *et al*., 2019). Similarly, no significant differences in growth were observed for the L-serine mutants or the *sdaABdsdA* triple mutant (Fig 8A). However, direct competition experiments revealed that loss of D-serine catabolism had no impact, loss of L-serine catabolism resulted in significant decrease in fitness compared to wild-type, and the combined loss of both D- and L-serine catabolism further decreased fitness (Fig 8B).

**Figure 8.**
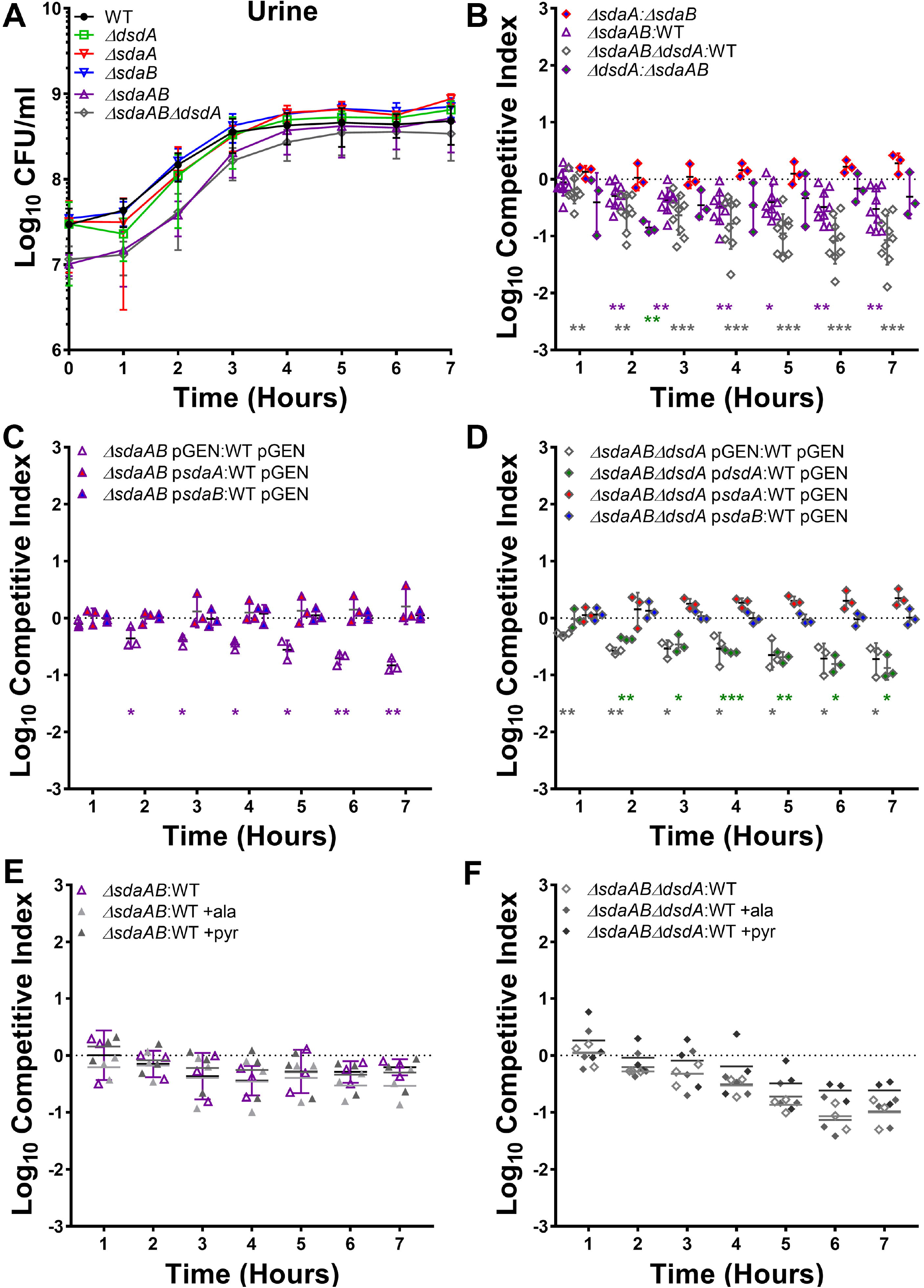
L-serine catabolism contributes to *P. mirabilis* fitness in human urine. (A) Wild-type *P. mirabilis* and indicated mutants were each cultured in 5 ml of filter-sterilized pooled human urine at 37°C with aeration, and growth was assessed by hourly dilution and plating for determination of CFUs/ml. Error bars represent mean ± SD from at least 3 independent experiments. (B-F) Urine was inoculated with a 50:50 mixture of the indicated strains and fitness was assessed by calculating a competitive index based on the CFUs of each strain at hourly time points. (C and D) Urine contained 100 µg/mL ampicillin to maintain plasmid selection. (E and F) Urine was supplemented with 10 mM alanine or pyruvate. Each data point indicates the log_10_ competitive index for the indicate mutant:WT or mutant:mutant inoculum from a single culture. Error bars represent the mean ± SD from at least 3 independent experiments, and the dashed line indicates Log_10_ CI=0 (the expected value if the ratio of mutant:WT is 1:1). \**p*<0.05, \*\**p*<0.01, \*\*\**p*<0.001 by Wilcoxon signed-rank test, with specific co-challenges indicated by color-coding.

Interestingly, competing the L-serine catabolism mutant against the D-serine catabolism mutant revealed only a slight fitness defect at a single time point (3 hours post-inoculation) (Fig 8B), suggesting that the ability to catabolize either D- or L-serine is sufficient for *P. mirabilis* fitness during growth in human urine and that neither provides a substantial fitness advantage over the other during this experimental timeframe. Similarly, no fitness defects were identified when competing the *sdaA* mutant against the *sdaB* mutant (Fig 8B), indicating that the presence of either L-serine deaminase is sufficient for fitness during growth in human urine. This was confirmed by complementation, as fitness of the *sdaAB* double mutant and the *sdaABdsdA* triple mutant could be fully restored by providing either *sdaA* or *sdaB* on a plasmid, while providing only *dsdA* to the triple mutant failed to restore fitness (Fig 8C and D). Supplementation with either alanine or pyruvate also failed to rescue the fitness defects of the double and triple mutant (Fig 8E and F), indicating that the fitness defects likely stem from alterations in membrane permeability rather than competition between serine and alanine during peptidoglycan synthesis or loss of energy production from serine.

### D- and L-serine catabolism only contribute to *P. mirabilis* biofilm development under nutrient-rich conditions

The ability to form persistent biofilm communities is an important aspect of *P. mirabilis* colonization and pathogenesis in the urinary tract, particularly in the context of catheterized individuals (Armbruster *et al*., 2018, White *et al*., 2021). We therefore sought to determine if serine utilization contributes to *P. mirabilis* biofilm formation. When biofilms were established in LB broth, loss of either D- or L-serine catabolism resulted in a significant decrease in biomass compared to wild-type *P. mirabilis* (Fig 9A). L-serine utilization appears to be the most critical factor as the Δ*sdaAB* biofilm defect was more pronounced than that of the Δ*dsdA* mutant and no further decrease was observed for the Δ*sdaAB*Δ*dsdA* mutant compared to Δ*sdaAB*. No differences in biofilm-associated CFUs were observed for any of the strains (Fig 9B), indicating that the biofilm defect is likely due to a decrease in production of matrix material or extracellular polymeric substances. Plasmid-based complementation experiments were not conducted due to issues with maintaining plasmid selection in *P. mirabilis* during biofilm formation. However, the defect of the Δ*dsdA* mutant was fully complemented by the addition of L-serine (Fig 9C), indicating that the defect for this mutant likely pertains to perturbation of L-serine biosynthesis as a result of intracellular accumulation of D-serine. In contrast, the Δ*sdaAB* and Δ*sdaAB*Δ*dsdA* mutants could not be complemented by the addition of either alanine or pyruvate, indicating that the biomass defects of L-serine catabolism mutants likely stems from disruption of membrane permeability rather than cell wall biosynthesis or energy production via catabolism to pyruvate (Fig 9D). Importantly, biofilm defects only occurred in LB, as no defects were observed for biofilms formed in human urine (Fig 9E).

**Figure 9.**
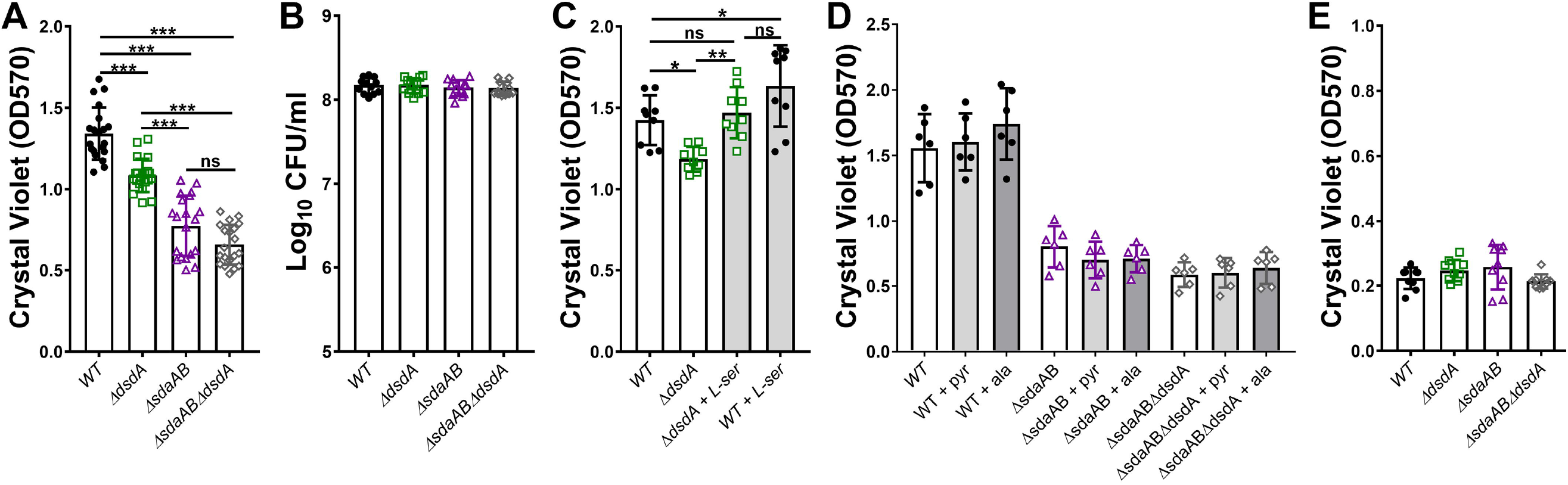
D- and L-serine catabolism both contribute to *P. mirabilis* biofilm formation in LB but not in human urine. Wild-type *P. mirabilis* and indicated mutants were suspended in either LB (A-D) or filter-sterilized pooled human urine (E) and distributed into 24-well plates. After a 20 hour incubated at 37°C under stationary conditions, biofilm biomass was quantified by crystal violet staining (A, C, D, and E) or by dilution and plating for determination of CFUs/ml (B). (C) LB was supplemented with 10 mM L-serine. (D) LB was supplemented with 10 mM alanine or pyruvate. Error bars represent mean ± SD for at least three independent experiments with at least 2 replicates each. \**p*<0.05, \*\**p*<0.01, \*\*\**p*<0.001 by one-way ANOVA with Dunnett’s test for multiple comparison. ns=not significant.

### Serine catabolism is critical for *P. mirabilis* fitness during CAUTI

We previously determined that D-serine catabolism provides *P. mirabilis* with a fitness advantage during experimental CAUTI (Brauer *et al*., 2019). To assess the contribution of L-serine catabolism to *P. mirabilis* colonization and survival within the catheterized urinary tract, female CBA/J mice were transurethrally inoculated with 10^5^ CFUs of either wild-type *P. mirabilis* HI4320, the *sdaAB* mutant, or the *sdaABdsdA* triple mutant. A 4 mm segment of sterile catheter tubing was placed in the bladder during inoculation, and mice were euthanized 96 hours post-inoculation. This infection model requires the bacteria to rapidly adapt to the urinary tract environment, including switching to utilizing peptides and amino acids as primary nutrients, establishing persistent biofilm communities on the catheter segment, and utilizing motility to ascend the ureters and infect the kidneys, all while resisting host innate immune defenses. Considering that many of these processes were impacted by loss of serine catabolism when tested *in vitro,* the *sdaAB* and *sdaABdsdA* mutants were expected to exhibit defects in persistence with the urine, colonization of the bladder, and ascension to the kidneys. Indeed, mice inoculated with the either the *sdaAB* double mutant or the *sdaABdsdA* triple mutant exhibited significantly reduced bacterial burden in the urine, bladder, and kidneys, indicating that loss of L-serine catabolism substantially impairs *P. mirabilis* colonization, ascension, and survival within the catheterized urinary tract (Fig 10A). A co-challenge strategy was also used to directly assess the relative fitness of each mutant compared to wild-type *P. mirabilis*. The *sdaAB* mutant was so highly outcompeted by wild-type *P. mirabilis* in all organs that it was only above the limit of detection in 2/10 mice after 96 hours (Fig 10B). Considering that the *sdaABdsdA* mutant had more severe fitness defects *in vitro,* the co-challenge with this mutant was only taken out to 48 hours instead of 96 hours. Even in this shortened timeframe, the triple mutant was completely outcompeted by wild-type *P. mirabilis* in all organs and was only above the limit of detection in 3/10 mice (Fig 10C). Together, these data underscore the critical importance of serine catabolism for *P. mirabilis* colonization and fitness within the urinary tract.

**Figure 10.**
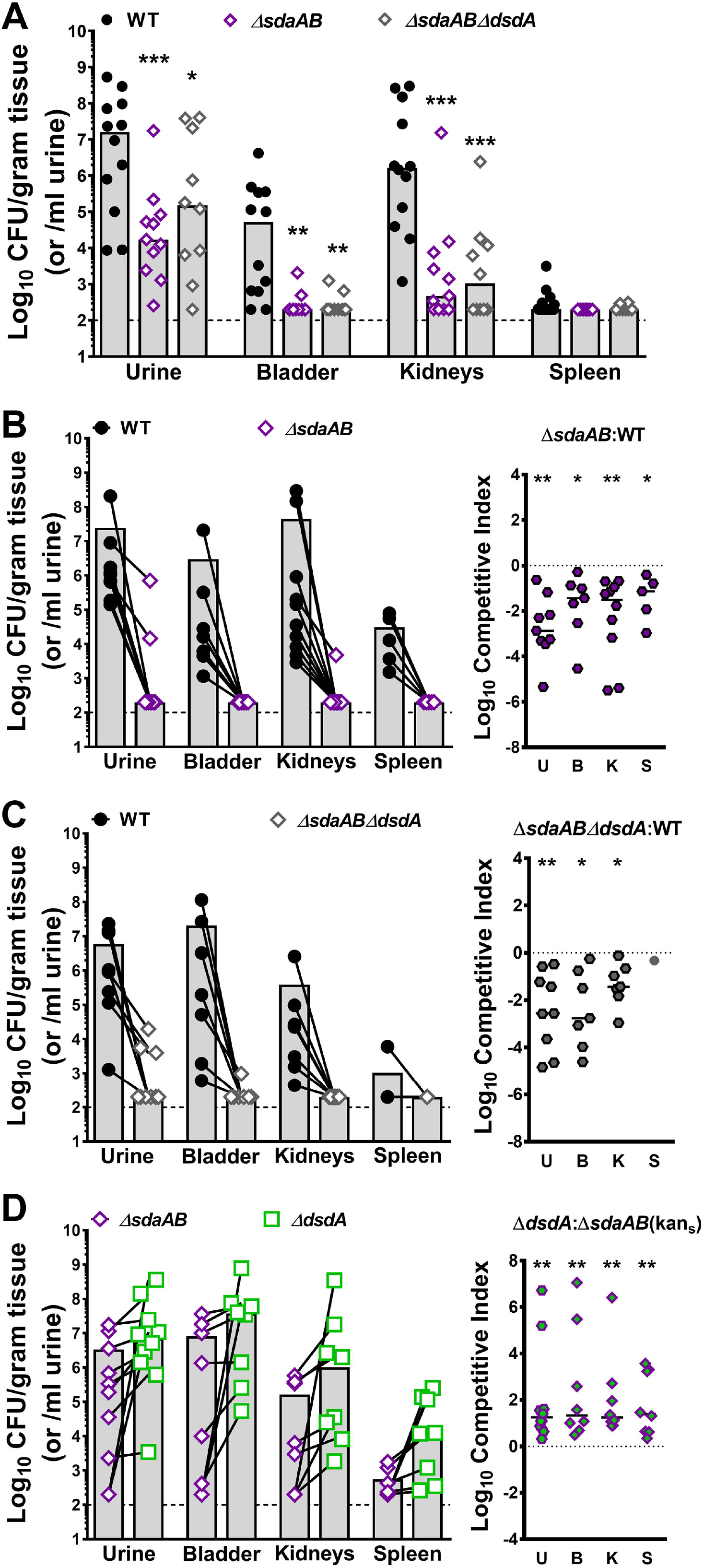
L-serine catabolism provides a greater fitness advantage to *P. mirabilis* within the catheterized urinary tract than D-serine catabolism. Wild-type *P. mirabilis,* the L-serine catabolism mutant (*sdaAB*) and the triple mutant (*sdaABdsdA*) were cultured in LB broth overnight, washed once in PBS, and adjusted to 2×10^6^ CFU/ml. (A) Female CBA/J mice were transurethrally inoculated with 50 µl containing 1×10^5^ CFU of either WT (n=12), *sdaAB* (n=11), or *sdaABdsdA* (n=10), and a 4 mm segment of silicone catheter tubing was advanced into the bladder during inoculation. Mice were euthanized 96 hours post-inoculation (hpi), and bacterial burden was determined in the urine, bladder, kidneys, and spleen. Each symbol represents the Log_10_ CFU per milliliter of urine or gram of tissue from an individual mouse, gray bars represent the median, and the dashed line indicates the limit of detection. *\*\*\*P*<0.01 by nonparametric Mann-Whitney test. (B-D) CBA/J mice were inoculated as above with the following 50:50 mixtures and euthanized at the indicated time points: (B) WT and *sdaAB* (n=10, 96 hpi); (C) WT and *sdaABdsdA* (n=10, 48 hpi); and (D) *dsdA* and *sdaAB* (n=10, 96 hpi). Each symbol represents the Log_10_ CFU per milliliter of urine or gram of tissue from an individual mouse, with CFUs of each strain recovered from an individual mouse connected by a black line. A competitive index was also calculated for each co-challenge, in which each symbol represents the Log_10_ CI for an individual mouse, error bars represent the median, and the dashed line indicates Log_10_ CI=0 (the expected value if the ratio of the indicated strains is 1:1). *\*P*<0.05, *\*\*P*<0.01 by Wilcoxon signed-rank test.

Notably, the serine utilization mutants both colonized all organs to a similar level during independent challenge, suggesting that the additional loss of D-serine catabolism had a negligible impact on colonization and survival within the urinary tract and that L-serine catabolism is the more critical factor for colonization of the catheterized urinary tract. We therefore sought to clarify the relative contribution of D-vs L-serine catabolism by directly competing the *dsdA* mutant against the *sdaAB* mutant (Fig 10D). In contrast to co-challenge with wild-type *P. mirabilis*, both mutants highly colonized all organs in the majority of mice. These data indicate that the relative fitness of these strains is much more similar to each other than to wild-type *P. mirabilis,* likely because they are no longer competing for import of their specific serine enantiomer. However, the *dsdA* mutant consistently achieved higher colonization levels than the *sdaAB* mutant in co-challenged mice, demonstrating that L-serine catabolism provides a greater overall advantage to *P. mirabilis* within the urinary tract than D-serine catabolism.

## Discussion

Amino acid import and catabolism have been considered critical for bacterial colonization and persistence within the urinary tract, yet few studies have examined amino acid utilization by uropathogens. In this study, we elucidated the hierarchy of amino acid import by *P. mirabilis* during growth in human urine, identified serine as the most abundant amino acid that is preferentially depleted by *P. mirabilis,* and explored the relative contribution of L- versus D-serine catabolism to pathogenic potential and fitness. The preferential depletion of L-serine followed by D-serine during growth in urine is particularly striking considering that serine was the 7th most abundant amino acid in this pool of normal human urine, while the concentration of the four most abundant amino acids either remained constant over time (valine, glycine, and alanine) or increased (methionine). This observation suggests that serine is either an important signaling factor for *P. mirabilis,* as has been recently proposed (Chittor & Gibbs, 2021), or that the specific catabolism of serine is important for rapid growth and fitness of *P. mirabilis* within urine and the urinary tract.

Prior studies found that disrupting serine catabolism can have deleterious impacts on growth due to perturbation of energy production, amino acid homeostasis, and cell wall biosynthesis due to intracellular serine accumulation (Anfora *et al*., 2007, Kriner & Subramaniam, 2020, Zhang *et al*., 2010, Zhang & Newman, 2008, Hama *et al*., 1991, Tazuya-Murayama *et al*., 2006). We therefore applied a chemical complementation strategy to assess the potential contribution of disrupted energy production and cell wall biosynthesis to each phenotype of the serine mutants (summarized in Table 2). This approach revealed that altered energy production via pyruvate is not a major contributor to the *in vitro* defects of serine catabolism mutants, and disruption of cell wall biosynthesis due to perturbation of the serine:alanine ratio only appears to impact motility. This approach further led to elucidation of a membrane permeability defect that likely stems from an alteration in membrane potential, which appears to be the underlying cause of the majority of *in vitro* defects for the serine catabolism mutants and is also present in serine import mutants. It is also notable that disruption of D-serine catabolism alone did not impact membrane permeability, and the majority of *in vitro* defects were specific to the triple mutant in which both D- and L-serine catabolism were disrupted. Thus, the majority of the defects for the serine catabolism mutants likely pertain to loss of serine utilization rather than intracellular serine accumulation.

**Table 2:** Summary of phenotypes associated with D vs L serine utilization.

| Phenotypes | $\Delta dsdA$ | $\Delta sdaAB$ | $\Delta sdaAB dsdA$ |
| --- | --- | --- | --- |
| Growth in PMSM | +/- | +/- | +/- |
| Ability to use D-serine as sole C or N source | --- | - | --- |
| Complementation: | *pdsdA* | NA | *pdsdA* |
| Ability to use L-serine as sole C or N source | +/- | --- | --- |
| Complementation: | NA | *psdaA* or *psdaB* | *psdaA* or *psdaB* |
| Growth in LB | +/- | +/- | +/- |
| Relative fitness in LB | +/- | - | -- |
| Complementation: | NA | Neither alanine nor pyruvate | Neither alanine nor pyruvate |
| Probable causes: | NA | Membrane potential, AA homeostasis | Membrane potential, AA homeostasis |
| Growth in urine | +/- | +/- | +/- |
| Relative fitness in urine | +/- | - | -- |
| Complementation: | NA | *psdaA* or *psdaB*, but not alanine or pyruvate | *psdaA* or *psdaB*, but not alanine or pyruvate |
| Probable causes: | NA | Membrane potential, AA homeostasis | Membrane potential, AA homeostasis |
| Antibiotic/detergent sensitivity | +/- | +/- | -- |
| Complementation: | NA | NA | Neither alanine nor pyruvate |
| Probable causes: | NA | Membrane potential, AA homeostasis | Membrane potential, AA homeostasis |
| Membrane permeability | +/- | -- | --- |
| Swarming motility | +/- | - | --- |
| Complementation: | NA | *psdaA*, *psdaB*, or alanine | Partially by *pdsdA*, fully by *psdaA*, *psdaB*, or alanine |
| Probable causes: | NA | Cell wall perturbation | Cell wall perturbation |
| Swimming motility | +/- | -- | --- |
| Complementation: | NA | Partially by alanine, fully by *psdaA* or *psdaB*, | Partially by *psdaB* or alanine, fully by *psdaA* |
| Probable causes: | NA | Cell wall perturbation, membrane potential, AA homeostasis | Cell wall perturbation, membrane potential, AA homeostasis |
| Biofilm formation in LB | - | -- | -- |
| Complementation: | L-serine | Neither alanine nor pyruvate | Neither alanine nor pyruvate |
| Probable causes: | Perturbation of L-serine synthesis | Membrane potential, AA homeostasis | Membrane potential, AA homeostasis |
| Biofilm formation in urine | +/- | +/- | +/- |
| CAUTI colonization | - | --- | --- |
| CAUTI relative fitness | --- | --- | --- |

(+/-) similar to wild-type; (-) slight defect; (--) moderate defect; (---) severe defect; NA not applicable

While our data clearly demonstrate that disrupting serine catabolism results in substantial colonization and fitness defects in the mouse model of CAUTI, we were not able to specifically determine whether the *in vivo* defects stemmed from loss of serine utilization or to the resulting alteration in membrane permeability and cell wall biosynthesis. Considering that none of the serine utilization mutants exhibited growth or biofilm defect when cultured individually in urine, and the majority of the observed defects pertained only to the triple mutant during growth in LB broth, we speculate that serine import and catabolism are the key factors underlying the colonization defect in the CAUTI model. Testing this hypothesis would require disrupting serine import and assessing colonization and fitness of the import mutant; however, our data demonstrate that *sdaC* and *sstT* contribute to import of both serine and threonine, and that at least one additional transporter is capable of importing serine. In *E. coli,* the YhaO inner membrane transporter was determined to be capable of importing either D- or L-serine (Connolly *et al*., 2016), and *P. mirabilis* strain HI4320 encodes a YhaO homolog (PMI2745). Fully disrupting both D- or L-serine import in HI4320 may therefore require mutation of *dsdX, sdaC, sstT,* and *yhaO,* and would also disrupt import of threonine. Future efforts are focused on dissecting the specific contributions of serine and threonine import and catabolism to *P. mirabilis* fitness within the urinary tract.

Our observation that the L-serine catabolism mutant and the D-serine catabolism mutant exhibited almost comparable fitness to each other during growth in urine *in vitro* with only a modest increase in fitness for the *dsdA* mutant in the mouse model of CAUTI was surprising, considering that the *dsdA* mutant exhibited fewer overall defects *in vitro* compared to the *sdaAB* mutant (summarized in Table 2). The differences in the *in vitro* defects of these mutants could be due to a lower overall level of D-serine accumulation in the *dsdA* mutant than L-serine accumulation in the *sdaAB* mutant when they are grown in LB broth, or to a more pronounced impact of L-serine accumulation on cell wall and membrane permeability. Alternatively, the ability to catabolize either enantiomer may be sufficient to support *P. mirabilis* colonization in an environment where they are present at a 1:1 ratio, such as during growth in urine. If so, L-serine catabolism would be expected to provide an even greater fitness advantage than D-serine catabolism in other niches, such as the bloodstream where the ratio of D- to L-serine is closer to 1:100 (Miyoshi *et al*., 2009, Bruckner & Schieber, 2001).

L-serine has previously been demonstrated to contribute to growth and survival of *Enterobacteriaceae,* especially in an inflammatory environment. For instance, *E. coli* typically prefers to catabolize sugars when available, but the environment of the inflamed gut induces a preferential catabolism of L-serine that is critical for *E. coli* expansion in a colitis model (Kitamoto *et al*., 2020). L-serine catabolism has also been demonstrated to contribute to intra-species competition and relative fitness in this niche, and may also provide an advantage in polymicrobial settings (Kitamoto *et al*., 2020). Considering that UTIs involving *P. mirabilis* are often polymicrobial (Gaston *et al*., 2021, Armbruster *et al*., 2017b), L-serine catabolism may become even more critical during coinfection. We previously demonstrated that D-serine catabolism remains an important fitness factor during coinfection with *Providencia stuartii* (Brauer et al., 2019), and L-serine catabolism genes *sdaA* and *sdaB* were also identified as candidate fitness factors for *P. mirabilis* CAUTI during coinfection with *P. stuartii* (Armbruster et al., 2017a). The contribution of these and other amino acid import and catabolism pathways to competition with other co-colonizers is an active area of further investigation. Ultimately, further elucidation of the nutrient preferences of common uropathogens could contribute to the development of dietary interventions aimed at reducing infection risk or disease severity.

## Experimental Procedures

### Ethics statement

Mouse infection studies were approved by the Institutional Animal Care and Use Committee (IACUC) at the State University of New York at Buffalo Jacobs School of Medicine and Biomedical Sciences (MIC31107Y), and were in accordance with the Office of Laboratory Animal Welfare (OLAW), the United States Department of Agriculture (USDA), and the guidelines specified by the Association for Assessment and Accreditation of Laboratory Animal Care, International (AAALAC, Intl.).

### Data availability statement

All data that support the findings of this study are available in the main text.

### Bacterial strains

*Proteus mirabilis* strain HI4320 was isolated from the urine of a catheterized nursing home resident (Mobley *et al*., 1985, Mobley & Warren, 1987). All mutants were generated by inserting a kanamycin resistance cassette into the gene of interest following the Sigma TargeTron group II intron protocol as previously described (Pearson & Mobley, 2007). The *P. mirabilis* L-serine degradation double mutant (Δ*sdaAB*) and D/L-serine degradation triple mutant (Δ*sdaAB*Δ*dsdA*) were constructed by following the TargeTron protocol for use with a *loxP* vector as previously described (Armbruster *et al*., 2013, Brauer *et al*., 2020). All mutants were verified by selection on kanamycin and by PCR.

Insertion of a kanamycin cassette via TargeTron retrohoming can result in polar effects. All mutants were therefore complemented by providing *dsdA*, *sdaA* or *sdaB* on a plasmid. Complementation vectors were constructed by amplifying *dsdA*, *sdaA* or *sdaB* along with approximately 500 bp of their flanking sequences and ligating the amplified sequences into a linearized pGEN-MCS vector. Successful complementation was verified by selection on ampicillin plates and by PCR. Primer sequences for generation and verification of L-serine mutants and for creating pGEN complementation vectors are provided in Table 3.

**Table 3:**
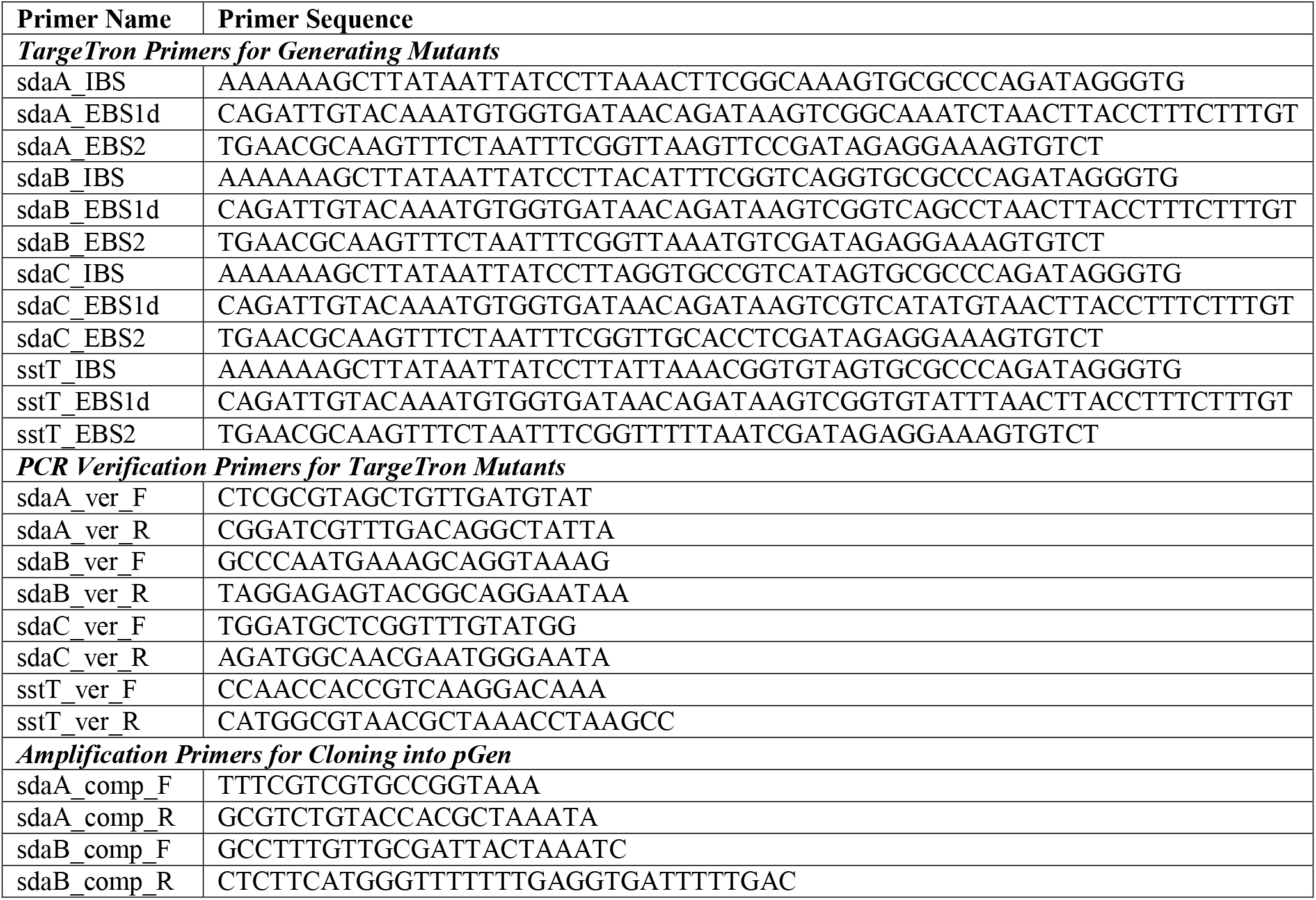
Primers used in this study.

### Culture conditions and media

Bacteria were routinely cultured at 37°C with shaking at 225 rpm in 5 ml low-salt LB broth (10 g/L tryptone, 5 g/L yeast extract, 0.1 g/L NaCl) or on low-salt LB solidified with 1.5% agar. *Proteus mirabilis* minimal salts medium (PMSM) (Belas *et al*., 1991) was used for studies requiring defined medium (10.5 g/L K_2_HPO_4_, 4.5 g/L KH_2_PO_4_, 1 g/L (NH_4_)_2_SO_4_, 15 g/L agar, supplemented with 0.001% nicotinic acid, 1mM MgSO_4_, and 0.2% glycerol). Where indicated, the PMSM formulation was modified by removing the glycerol or ammonium sulfate and replacing with 10 mM D-serine, L-serine, or L-threonine. Filter-sterilized pooled human urine from at least 20 de-identified female donors was purchased from Cone Bioproducts (Seguin, TX) and stored at −80°C for use as a physiologically-relevant growth medium. Media were supplemented with kanamycin (50 µg/mL) or ampicillin (100 µg/mL) as needed for selection, as well as 10 mM L-alanine, 10 mM pyruvate, or 10 mM pantothenate as indicated.

### Quantification of amino acids by HPLC

Pooled human urine was either left uninoculated or inoculated with 10^7^ CFU/ml of *P. mirabilis* strain HI4320 and incubated at 37°C with shaking at 225 rpm. Immediately upon inoculation and every 30 minutes thereafter for 12 hours, a 1 mL aliquot was removed, centrifuged at 10,000xg for 3 min, then filtered through a 0.22 µm Spin-s column (Corning 8160, Corning,NY) at 10,000xg for 3 min. Filtrate was then frozen and stored at −80□C.

The following solvents were prepared according to Agilent’s Amino Acid Analysis “How-To” Guide (https://www.agilent.com/cs/library/brochures/5991-7694EN_AdvanceBio%20AAA_How-To%20Guide_LR.pdf): Mobile Phase A (10 mM Na_2_HPo_4_ + 10mM Na_2_B_4_o_7_ in mili-Q water); Mobile Phase B (45:45:10 Acetonitrile:Methanol: mili-Q water); Injection diluent (100 mL Mobile phase A + 0.4 mL concentrated Phosphoric acid). All other reagents came from the AdvanceBio amino acid kit (5190-9426).

The HPLC setup included an Agilent G7129A 1260 Vial Sampler, an Agilent G7111A Quaternary Pump, and an Agilent G7121A 1260 FLD. Samples were run on an AdvanceBio AAA column (PN 655950-802) with a UPLC guard (PN 820750-931). A column heater is recommended by Agilent but was not available, so samples were run at room temperature. Prior to each run, frozen urine supernatants were brought up to room temperature and vortexed to re-dissolve any precipitation. 20 µL of urine supernatant was added to 180 µl of sample diluent (100 µM Norvaline prepared in Injection diluent) in Agilent 250 µL polypropylene vial inserts. Urine and sample diluent were gently mixed by hand to avoid formation of air bubbles in the vial inserts. Run parameters were modified from Agilent’s Amino Acid Analysis “How-To” Guide. Due to pressure limitations in our system (maximum 400 Bar), the protocol was modified as follows: the flow rate was set to 1.2 ml/min; the time of gradient from 43-100% of mobile phase B was increased to 29 min 45 sec; the washout time was increased for a total run time of 33 min/sample. In order to simplify the protocol, we removed the FMOC derivatization of proline and hydroxyproline by substituting a second vial of borate buffer (PN 5061-3335) in the online derivatization.

Peaks were called as specific amino acids based on their retention times in comparison to an Agilent amino acid standard (PN 5061-3330) spiked with asparagine, glutamine, and tryptophan, as detailed in Agilent’s Amino Acid Analysis “How-To” Guide. The area under the curve for all amino acids in a given sample were normalized to norvaline as an internal control for run-to-run and loading variability. Amino acid concentrations were determined via interpolation from a standard curves, which was generated using the Agilent amino acid standard spiked with asparagine, glutamine, and tryptophan.

### D-serine and L-serine measurement in urine

Total serine concentration and assessment of D- vs L- enantiomers was conducted as previously described using a fluorometric DL-Serine Assay Kit (BioVision) (Brauer *et al*., 2019). Briefly, aliquots generated from the above HPLC experiments were deproteinized using the sample cleanup mix provided in the kit, filtered through a 10 kDa molecular weight cutoff spin column, and assayed in triplicate following the manufacturer protocol.

### Growth curves

Overnight cultures of wild-type *P. mirabilis* HI4320 or serine utilization mutants were adjusted to OD_600_ of 0.02 (∼2×10^7^ CFU/ml) in various formulations of PMSM, LB, or urine. For assessment of growth by optical density, 200 µl of bacterial suspensions were distributed into at least 3 wells each of a clear 96-well plate and incubated at 37°C with continuous double-orbital shaking in a BioTek Synergy H1 96-well plate reader, with a 1°C temperature differential between the top and bottom of the plate to prevent condensation. Bacterial growth was assessed by absorbance (OD_600_) at 15 min intervals for a duration of 18 hours. For assessment by CFUs, 5 ml bacterial suspensions were incubated at 37°C with shaking at 225 rpm and aliquots were taken hourly for serial dilution and plating onto LB agar using an EddyJet 2 spiral plater (Neutec Group) for determination of CFUs using a ProtoCOL 3 automated colony counter (Synbiosis).

### Co-culture competition experiments

The relative fitness of serine utilization mutants was assessed by direct competition with either wild-type *P. mirabilis* HI4320 or other mutants as indicated. PMSM, LB, or urine were inoculated with a 1:1 mixture of the strains of interest and incubated at 37°C with shaking at 225 rpm. Aliquots were taken hourly for determination of bacterial CFUs by plating onto LB agar (total CFUs) and LB with kanamycin (CFUs of the mutant strain). A competitive index (CI) was calculated as follows:

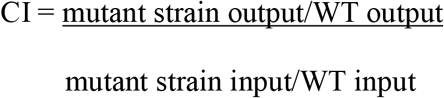

Log_10_CI=0 indicates that the ratio of the strains in the output is similar to the input, and neither strain had an advantage, Log_10_CI>0 indicates that the mutant has a competitive advantage over WT, and Log_10_CI<0 indicates that the mutant is outcompeted by WT.

### Disc diffusion assays

Overnight broth cultures of bacterial strains of interest were used to generate bacterial lawns by swabbing LB agar plates in three different directions. Swabbed plates were briefly air dried, then antibiotic or detergent discs were placed in triplicate using sterile forceps. Discs containing 10µg imipenem, 5µg ciproflxacin or 30µg ceftazidime were purchased from Hardy Diagnostics, while detergent discs were made by soaking blank paper discs (Becton Dickinson) for 5 minutes at room temperature in individual wells of a 24-well tissue culture plate filled with 500 µL 20% sodium dodecyl sulfate or 20% sodium deoxycholate (Sigma). After disc placement, plates were incubated at 37°C for 18 hours and zones of inhibition were measured for using EZCal digital calipers.

### Membrane permeability assay

1-*N*-phenylnaphthylamine (NPN) uptake and fluorescence was used to assess membrane permeability as described elsewhere (Helander & Mattila-Sandholm, 2000). Briefly, overnight cultures of wild-type *P. mirabilis* or indicated mutants were diluted 1:100 in 20 mL LB and incubated at 37°C with shaking at 225 rpm to mid-log phase. When cultures reached an OD_600_ of 0.5, 10 mL of culture was centrifuged at 3000 x g for 20 minutes, the supernatant was removed, and the cell pellet was resuspended in 5 mL of 5 mM 4-(2- hydroxyethyl)-1-piperazineethanesulfonic acid (HEPES) buffer at pH 7.2. 100 μl of the bacterial suspension, 50 μl of 40 μmol/L NPN prepared in HEPES, 25 μl of 0.5M ethylenediaminetetraacetic acid (EDTA) at pH 8.0, and HEPES buffer were then added as indicated to triplicate wells of a Corning Incorporated (Costar^R^) 96 well black clear bottom microtiter plate to a final assay volume of 200 μl. Fluorescence was immediately measured at 355 nm excitation and 405 nm emission using a BioTek Synergy H1 plate reader. Each assay was performed in triplicate, and the NPN uptake factor was calculated as follows:

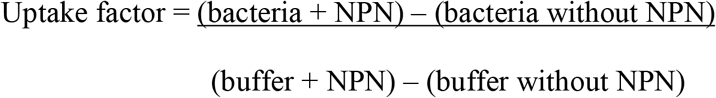

### Acid tolerance assay

Overnight cultures of strains of interest were diluted 1:50 in LB and incubated at 37°C with aeration for 2-3 hours until log phase. Culture were then centrifuged and resuspended in 10 mM 2-(*N*-morpholino)-ethanesulfonic acid (MES; Sigma) buffer to an OD_600_ of ∼ 1.0. The bacterial suspension was then diluted 1:10 in 1 ml of 10 mM MES adjusted to pH 2.5, 5, or 7 with or without 10mM D- or L-serine. Samples were incubated at 37°C with aeration for 1 hour and plated using an EddyJet 2 spiral plater (Neutec Group) for determination of CFUs using a ProtoCOL 3 automated colony counter (Synbiosis).

### Motility

Swarming motility was assessed using swarm agar (LB agar with 5 g/L NaCl). Briefly, strains of interest were incubated overnight in LB broth and 5 µl was spotted onto the center of at least 3 replicate swarming plates per experiment. Plates were then incubated at 37°C for ∼18 hours and swarm diameter was measured. Swimming motility was assessed using Mot medium (10 g/L tryptone and 5 g/L NaCl solidified with 0.3% agar). Briefly, overnight cultures of strains of interest were stab-inoculated into the center of the agar plate. Plates were then incubated at 30°C for ∼18 hours without inverting, and swim diameter was measured. Where indicated, swim and swarm media were supplemented with ampicillin (100 µg/mL) to maintain selection of complementation vectors or 10 mM L-alanine, 10 mM pyruvate, or 10 mM pantothenate to attempt to rescue the serine utilization mutants.

### Biofilm assays

Overnight cultures of strains of interest were adjusted to ∼2 x 10^7^ CFU/ml in LB broth and 1.5 mL was dispensed into triplicate wells of tissue culture treated 24 well plates (Falcon 353047). Uninoculated sterile LB was also dispensed into triplicate wells to serve as a blank for background crystal violet staining. Plates were incubated for 20 hours in a stationary incubator at 37°C, after which time supernatants were carefully removed and adherent biofilms were air dried for five minutes. Biofilms were then stained with 1.5 mL 0.1% crystal violet for 10 minutes, washed once with 1.5 mL distilled water, and solubilized in 2 mL 95% ethanol for 20 minutes on an orbital shaker at 200 rpm. A micropipette tip was used to scrape the bottom and sides of the wells to fully resuspend all stained biomass. Crystal violet absorbance was measured at 570 nm using a BioTek Synergy H1 plate reader.

### Mouse model of CAUTI

CAUTI studies were carried out as previously described (Armbruster *et al*., 2017a, Brauer *et al*., 2019). Briefly, the inoculum was prepared by washing overnight cultures of strains of interest in phosphate-buffered saline (PBS: 0.128 M NaCl, 0.0027 M KCl, pH 7.4), adjusting to OD_600_ 0.2 (∼2×10^8^ CFU/ml), and diluting 1:100 to achieve an inoculum of 2×10^6^ CFU/ml. Female CBA/J mice aged 6-8 weeks (Jackson Laboratory) were anesthetized with a weight-appropriate dose (0.1 ml for a mouse weighing 20 gm) of ketamine/xylazine (80- 120 mg/kg ketamine and 5-10 mg/kg xylazine) by IP injection and inoculated transurethrally with 50 µl of the diluted suspension (1×10^5^ CFU/mouse), and a 4 mm segment of sterile silicone tubing (0.64 mm O.D., 0.30 mm I.D., Braintree Scientific, Inc.) was carefully advanced into the bladder during inoculation and retained for the duration of the study as described previously (Guiton *et al*., 2010, Armbruster *et al*., 2017c). After 96 hours, urine was collected, mice were euthanized, and bladders, kidneys, and spleens were harvested into 5 mL Eppendorf tubes containing 1 mL PBS. Tissues were homogenized using a Bullet Blender 5 Gold (Next Advance) and plated using an EddyJet 2 spiral plater (Neutec Group) for determination of CFUs using a ProtoCOL 3 automated colony counter (Synbiosis). For co-challenge experiments, mice were inoculated with a 50:50 mix of strains of interest and euthanized at either 48 hours or 96 hours as indicated, and samples were plated onto plain LB agar (total CFUs) and LB with kanamycin (mutant strain CFUs), and a competitive index was calculated as described above.

### Statistical analysis

For motility, biofilm, disc diffusion, growth curves, and CFU data, significance was assessed using two-way analysis of variance (ANOVA) corrected for multiple comparisons. Competitive indices from co-challenge experiments were assessed by Wilcoxon signed-rank test. These analyses were performed using GraphPad Prism, version 7.03 (GraphPad Software). All *P* values are two tailed at a 95% confidence interval.

## Acknowledgements

We would like to thank Dr. Thomas Russo and members of his laboratory for helpful comments and critiques, as well as Dr. Michael Malkowski and members of his laboratory for their assistance with HPLC. This work was funded by the National Institute of Diabetes and Digestive and Kidney Diseases under award R01 DK123158 to CEA. The content is solely the responsibility of the authors and does not necessarily represent the official views of the National Institutes of Health.

## Author contributions

Study design: CEA

Data acquisition, analysis, and interpretation: ALB, BSL, SMT, ND, BCH, CEA

Writing and editing of the manuscript: CEA, ALB, BSL, SMT, ND, BCH

## References

(2021) Mayo Clinic Laboratories Test Catalog: AAPD. In: Amino Acids, Quantitative, Random, Urine. pp.

Anfora, A.T., Haugen, B.J., Roesch, P., Redford, P., and Welch, R.A. (2007) Roles of serine accumulation and catabolism in the colonization of the murine urinary tract by Escherichia coli CFT073. Infection and immunity 75: 5298–5304.

Armbruster, C., Mobley, H., and Pearson, M. (2018) Pathogenesis of *Proteus mirabilis* Infection. EcoSal Plus.

Armbruster, C.E., Brauer, A.L., Humby, M.S., Shao, J., and Chakraborty, S. (2021) Prospective assessment of catheter-associated bacteriuria clinical presentation, epidemiology, and colonization dynamics in nursing home residents. JCI Insight 6: 2020.2009.2029.20204107.

Armbruster, C.E., Forsyth-DeOrnellas, V., Johnson, A.O., Smith, S.N., Zhao, L., Wu, W., and Mobley, H.L.T. (2017a) Genome-wide transposon mutagenesis of *Proteus mirabilis*: Essential genes, fitness factors for catheter-associated urinary tract infection, and the impact of polymicrobial infection on fitness requirements. PLOS Pathogens 13: e1006434.

Armbruster, C.E., Forsyth, V.S., Johnson, A.O., Smith, S.N., White, A.N., Brauer, A.L., Learman, B.S., Zhao, L., Wu, W., Anderson, M.T., Bachman, M.A., and Mobley, H.L.T. (2019) Twin arginine translocation, ammonia incorporation, and polyamine biosynthesis are crucial for *Proteus mirabilis* fitness during bloodstream infection. PLoS Pathog 15: e1007653.

Armbruster, C.E., Hodges, S.A., and Mobley, H.L. (2013) Initiation of swarming motility by Proteus mirabilis occurs in response to specific cues present in urine and requires excess L-glutamine. J Bacteriol 195: 1305–1319.

Armbruster, C.E., Hodges, S.A., Smith, S.N., Alteri, C.J., and Mobley, H.L. (2014) Arginine promotes *Proteus mirabilis* motility and fitness by contributing to conservation of the proton gradient and proton motive force. MicrobiologyOpen 3: 630–641.

Armbruster, C.E., Prenovost, K., Mobley, H.L.T., and Mody, L. (2017b) How Often Do Clinically Diagnosed Catheter-Associated Urinary Tract Infections in Nursing Home Residents Meet Standardized Criteria? J Am Geriatr Soc 65: 395–401.

Armbruster, C.E., Smith, S.N., Johnson, A.O., DeOrnellas, V., Eaton, K.A., Yep, A., Mody, L., Wu, W., and Mobley, H.L.T. (2017c) The Pathogenic Potential of *Proteus mirabilis* is Enhanced by Other Uropathogens During Polymicrobial Urinary Tract Infection. Infection and Immunity 85: e00808–00816.

Belas, R., Erskine, D., and Flaherty, D. (1991) Transposon mutagenesis in *Proteus mirabilis*. J Bacteriol 173: 6289–6293.

Bouatra, S., Aziat, F., Mandal, R., Guo, A.C., Wilson, M.R., Knox, C., Bjorndahl, T.C., Krishnamurthy, R., Saleem, F., Liu, P., Dame, Z.T., Poelzer, J., Huynh, J., Yallou, F.S., Psychogios, N., Dong, E., Bogumil, R., Roehring, C., and Wishart, D.S. (2013) The Human Urine Metabolome. PLOS ONE 8: e73076.

Brauer, A.L., Learman, B.S., and Armbruster, C.E. (2020) Ynt is the primary nickel import system used by *Proteus mirabilis* and specifically contributes to fitness by supplying nickel for urease activity. Mol Microbiol 114: 185–199.

Brauer, A.L., White, A.N., Learman, B.S., Johnson, A.O., and Armbruster, C.E. (2019) D-Serine degradation by *Proteus mirabilis* contributes to fitness during single-species and polymicrobial catheter-associated urinary tract infection. mSphere 4.

Brooks, T., and Keevil, C.W. (1997) A simple artificial urine for the growth of urinary pathogens. Lett Appl Microbiol 24: 203–206.

Bruckner, H., and Schieber, A. (2001) Determination of amino acid enantiomers in human urine and blood serum by gas chromatography-mass spectrometry. Biomed Chromatogr 15: 166–172.

Burall, L.S., Harro, J.M., Li, X., Lockatell, C.V., Himpsl, S.D., Hebel, J.R., Johnson, D.E., and Mobley, H.L.T. (2004) *Proteus mirabilis* Genes That Contribute to Pathogenesis of Urinary Tract Infection: Identification of 25 Signature-Tagged Mutants Attenuated at Least 100-Fold. Infect. Immun. 72: 2922–2938.

Chittor, A., and Gibbs, K.A. (2021) The conserved serine transporter SdaC moonlights to enable self recognition. J Bacteriol: JB0034721.

Colomer-Winter, C., Flores-Mireles, A.L., Kundra, S., Hultgren, S.J., Lemos, J.A., and Dunman, P. (2019) (p)ppGpp and CodY Promote Enterococcus faecalis Virulence in a Murine Model of Catheter-Associated Urinary Tract Infection. mSphere 4: e00392–00319.

Connolly, J.P.R., Gabrielsen, M., Goldstone, R.J., Grinter, R., Wang, D., Cogdell, R.J., Walker, D., Smith, D.G.E., and Roe, A.J. (2016) A Highly Conserved Bacterial D-Serine Uptake System Links Host Metabolism and Virulence. PLoS pathogens 12: e1005359–e1005359.

Cosloy, S.D., and McFall, E. (1973) Metabolism of S-Serine in Escherichia coli K-12: Mechanism of Growth Inhibition. Journal of Bacteriology 114: 685–694.

Dunstan, R.H., Sparkes, D.L., Macdonald, M.M., De Jonge, X.J., Dascombe, B.J., Gottfries, J., Gottfries, C.G., and Roberts, T.K. (2017) Diverse characteristics of the urinary excretion of amino acids in humans and the use of amino acid supplementation to reduce fatigue and sub-health in adults. Nutrition Journal 16: 19.

Forsyth, V.S., Armbruster, C.E., Smith, S.N., Pirani, A., Springman, A.C., Walters, M.S., Nielubowicz, G.R., Himpsl, S.D., Snitkin, E.S., and Mobley, H.L.T. (2018) Rapid Growth of Uropathogenic Escherichia coli during Human Urinary Tract Infection. mBio 9.

Foxman, B., and Brown, P. (2003) Epidemiology of urinary tract infections: transmission and risk factors, incidence, and costs. Infect Dis Clin North Am 17: 227–241.

Gaston, J.R., Johnson, A.O., Bair, K.L., White, A.N., and Armbruster, C.E. (2021) Polymicrobial interactions in the urinary tract: is the enemy of my enemy my friend? Infect Immun.

Gibreel, T.M., Dodgson, A.R., Cheesbrough, J., Bolton, F.J., Fox, A.J., and Upton, M. (2012) High metabolic potential may contribute to the success of ST131 uropathogenic Escherichia coli. J Clin Microbiol 50: 3202–3207.

Gordon, D.M., and Riley, M.A. (1992) A theoretical and experimental analysis of bacterial growth in the bladder. Mol Microbiol 6: 555–562.

Griffith, D.P., Musher, D.M., and Itin, C. (1976) Urease. The primary cause of infection-induced urinary stones. Invest Urol 13: 346–350.

Guiton, P.S., Hung, C.S., Hancock, L.E., Caparon, M.G., and Hultgren, S.J. (2010) Enterococcal Biofilm Formation and Virulence in an Optimized Murine Model of Foreign Body-Associated Urinary Tract Infections. Infection and Immunity 78: 4166–4175.

Hama, H., Kayahara, T., Tsuda, M., and Tsuchiya, T. (1991) Inhibition of homoserine dehydrogenase I by L-serine in *Escherichia coli*. Journal of biochemistry 109: 604–608.

Helander, I.M., and Mattila-Sandholm, T. (2000) Fluorometric assessment of Gram-negative bacterial permeabilization. Journal of Applied Microbiology 88: 213–219.

Heßlinger, C., Fairhurst, S.A., and Sawers, G. (1998) Novel keto acid formate-lyase and propionate kinase enzymes are components of an anaerobic pathway in Escherichia coli that degrades L-threonine to propionate. Molecular Microbiology 27: 477–492.

Himpsl, S.D., Lockatell, C.V., Hebel, J.R., Johnson, D.E., and Mobley, H.L.T. (2008) Identification of virulence determinants in uropathogenic *Proteus mirabilis* using signature-tagged mutagenesis. J Med Microbiol 57: 1068–1078.

Johnson, A.O., Forsyth, V., Smith, S.N., Learman, B.S., Brauer, A.L., White, A.N., Zhao, L., Wu, W., Mobley, H.L.T., and Armbruster, C.E. (2020) Transposon Insertion Site Sequencing of *Providencia stuartii*: Essential Genes, Fitness Factors for Catheter-Associated Urinary Tract Infection, and the Impact of Polymicrobial Infection on Fitness Requirements. mSphere 5.

Kim, B.N., Kim, N.J., Kim, M.N., Kim, Y.S., Woo, J.H., and Ryu, J. (2003) Bacteraemia due to tribe Proteeae: a review of 132 cases during a decade (1991-2000). Scand J Infect Dis 35: 98–103.

Kitamoto, S., Alteri, C.J., Rodrigues, M., Nagao-Kitamoto, H., Sugihara, K., Himpsl, S.D., Bazzi, M., Miyoshi, M., Nishioka, T., Hayashi, A., Morhardt, T.L., Kuffa, P., Grasberger, H., El-Zaatari, M., Bishu, S., Ishii, C., Hirayama, A., Eaton, K.A., Dogan, B., Simpson, K.W., Inohara, N., Mobley, H.L.T., Kao, J.Y., Fukuda, S., Barnich, N., and Kamada, N. (2020) Dietary l-serine confers a competitive fitness advantage to Enterobacteriaceae in the inflamed gut. Nature Microbiology 5: 116–125.

Korte-Berwanger, M., Sakinc, T., Kline, K., Nielsen, H.V., Hultgren, S., and Gatermann, S.G. (2013) Significance of the D-Serine-Deaminase and D-Serine Metabolism of Staphylococcus saprophyticus for Virulence. Infection and Immunity 81: 4525–4533.

Kriner, M.A., and Subramaniam, A.R. (2020) The serine transporter SdaC prevents cell lysis upon glucose depletion in *Escherichia coli*. MicrobiologyOpen 9: e960.

Li, X., Zhao, H., Lockatell, C.V., Drachenberg, C.B., Johnson, D.E., and Mobley, H.L.T. (2002) Visualization of *Proteus mirabilis* within the Matrix of Urease-Induced Bladder Stones during Experimental Urinary Tract Infection. Infect. Immun. 70: 389–394.

Miyoshi, Y., Hamase, K., Tojo, Y., Mita, M., Konno, R., and Zaitsu, K. (2009) Determination of d-serine and d-alanine in the tissues and physiological fluids of mice with various d-amino-acid oxidase activities using two-dimensional high-performance liquid chromatography with fluorescence detection. Journal of Chromatography B 877: 2506–2512.

Mobley, H.L., Chippendale, G.R., Fraiman, M.H., Tenney, J.H., and Warren, J.W. (1985) Variable phenotypes of *Providencia stuartii* due to plasmid-encoded traits. J Clin Microbiol 22: 851–853.

Mobley, H.L.T., and Warren, J.W. (1987) Urease-Positive Bacteriuria and Obstruction of Long-Term Urinary Catheters. J Clin Microbiol 25: 2216–2217.

Nørregaard-Madsen, M., McFall, E., and Valentin-Hansen, P. (1995) Organization and transcriptional regulation of the Escherichia coli K-12 D-serine tolerance locus. Journal of Bacteriology 177: 6456–6461.

Parveen, S., and Reddy, M. (2017) Identification of YfiH (PgeF) as a factor contributing to the maintenance of bacterial peptidoglycan composition. Mol Microbiol 105: 705–720.

Paudel, S., Bagale, K., Patel, S., Kooyers, N.J., Kulkarni, R., and Dozois, C.M. (2021) Human Urine Alters Methicillin-Resistant Staphylococcus aureus Virulence and Transcriptome. Applied and environmental microbiology 87: e00744–00721.

Pearson, M.M., and Mobley, H.L.T. (2007) The type III secretion system of *Proteus mirabilis* HI4320 does not contribute to virulence in the mouse model of ascending urinary tract infection. J Med Microbiol 56: 1277–1283.

Pearson, M.M., Yep, A., Smith, S.N., and Mobley, H.L.T. (2011) Transcriptome of *Proteus mirabilis* in the Murine Urinary Tract: Virulence and Nitrogen Assimilation Gene Expression. Infect. Immun.: IAI.05152–05111.

Reitzer, L., and Zimmern, P. (2019) Rapid Growth and Metabolism of Uropathogenic Escherichia coli in Relation to Urine Composition. Clinical microbiology reviews 33: e00101–00119.

Sabih, A., and Leslie, S.W., (2022) Complicated Urinary Tract Infections. In: StatPearls. Treasure Island (FL), pp.

Sakinç, T., Michalski, N., Kleine, B., and Gatermann, S.G. (2009) The uropathogenic species *Staphylococcus saprophyticus* tolerates a high concentration of d-serine. FEMS Microbiology Letters 299: 60–64.

Schreiber, H.L.t., Conover, M.S., Chou, W.-C., Hibbing, M.E., Manson, A.L., Dodson, K.W., Hannan, T.J., Roberts, P.L., Stapleton, A.E., Hooton, T.M., Livny, J., Earl, A.M., and Hultgren, S.J. (2017) Bacterial virulence phenotypes of Escherichia coli and host susceptibility determine risk for urinary tract infections. Science translational medicine 9: eaaf1283.

Sintsova, A., Frick-Cheng, A.E., Smith, S., Pirani, A., Subashchandrabose, S., Snitkin, E.S., and Mobley, H. (2019) Genetically diverse uropathogenic Escherichia coli adopt a common transcriptional program in patients with UTIs. eLife 8: e49748.

Tazuya-Murayama, K., Aramaki, H., Mishima, M., Saito, K., Ishida, S., and Yamada, K. (2006) Effect of L-serine on the biosynthesis of aromatic amino acids in *Escherichia coli*. J Nutr Sci Vitaminol 52: 256–260.

Tielen, P., Rosin, N., Meyer, A.-K., Dohnt, K., Haddad, I., Jänsch, L., Klein, J., Narten, M., Pommerenke, C., Scheer, M., Schobert, M., Schomburg, D., Thielen, B., and Jahn, D. (2013) Regulatory and Metabolic Networks for the Adaptation of Pseudomonas aeruginosa Biofilms to Urinary Tract-Like Conditions. PLOS ONE 8: e71845.

Wagenlehner, F.M.E., Bjerklund Johansen, T.E., Cai, T., Koves, B., Kranz, J., Pilatz, A., and Tandogdu, Z. (2020) Epidemiology, definition and treatment of complicated urinary tract infections. Nat Rev Urol 17: 586–600.

Watanakunakorn, C., and Perni, S.C. (1994) *Proteus mirabilis* bacteremia: a review of 176 cases during 1980-1992. Scand J Infect Dis 26: 361–367.

White, A.N., Learman, B.S., Brauer, A.L., and Armbruster, C.E. (2021) Catalase Activity is Critical for *Proteus mirabilis* Biofilm Development, Extracellular Polymeric Substance Composition, and Dissemination during Catheter-Associated Urinary Tract Infection. Infect Immun 89: e0017721.

Zhang, X., El-Hajj, Z.W., and Newman, E. (2010) Deficiency in L-serine deaminase interferes with one-carbon metabolism and cell wall synthesis in *Escherichia coli* K-12. J Bacteriol 192: 5515–5525.

Zhang, X., and Newman, E. (2008) Deficiency in l-serine deaminase results in abnormal growth and cell division of *Escherichia coli* K-12. Molecular Microbiology 69: 870–881.

Zhao, H., Li, X., Johnson, D.E., and Mobley, H.L.T. (1999) Identification of protease and *rpoN*-associated genes of uropathogenic *Proteus mirabilis* by negative selection in a mouse model of ascending urinary tract infection. Microbiology 145: 185–195.

